# Mapping vascular plasticity in liver fibrogenesis identifies novel fibrosis-associated endothelial cells in early-stage liver disease

**DOI:** 10.64898/2026.03.12.710459

**Authors:** Christina Gkantsinikoudi, Joshua P Dignam, Raju Kumar, Elliot Jokl, Wenhao Li, Maryna Samus, Stephanie Landi, Varinder S Athwal, Timothy J Kendall, Antal Rot, Jonathan A Fallowfield, Karen Piper Hanley, William Alazawi, Neil P Dufton

## Abstract

Vascular plasticity is a crucial biological asset enabling our bodies to rapidly adapt to infections and acute inflammation. However, repeated insult during chronic disease can result in these vascular adaptations becoming irreversible, thereby driving disease progression and fibrosis. This study aimed to understand if phenotypic changes in endothelial cell (EC) identity could be indicative of progressive fibrosis and thereby offer new diagnostic and therapeutic opportunities for patients with metabolic dysfunction-associated steatotic liver disease (MASLD). Previous research has documented that a significant shift in EC transcriptomic signature occurs during liver fibrosis in both pre-clinical models and patients. However, the protein expression profile, phenotype and functional role of these new EC subpopulations that are induced during fibrogenesis is unclear. In this study, we integrate high-resolution imaging, proteomic and transcriptomic analysis which collectively highlight a central role for endothelial-to-mesenchymal transition (EndMT)-induced EC plasticity in the derivation of ‘fibrosis-associated’ EC (FAEC). We demonstrate that: 1) full spectrum flow cytometry can provide new opportunities to categorize and phenotype EC subpopulations, 2) two distinct EndMT-derived FAEC subpopulations expand during fibrogenesis; THY1.2^+^ICAM1^+^ and TAGLN^+^MCAM^+^ EC that display unique immunomodulatory and metabolic phenotypes, 3) TAGLN^+^ FAEC are a conserved, pro-fibrotic cell type arising at early stages of MASLD, and 4) increased hepatic expression of TAGLN is significantly associated with detrimental patient outcomes at all stages of liver disease. This study will pave the way for the development of FAEC-specific diagnostic and therapeutic approaches to tackle progressive fibrotic disease.

**One Sentence Summary:** Phenotyping fibrosis-associated endothelial cells reveals pro-fibrotic and immunomodulatory subpopulations at early stages of liver disease.

**Graphical Abstract:** 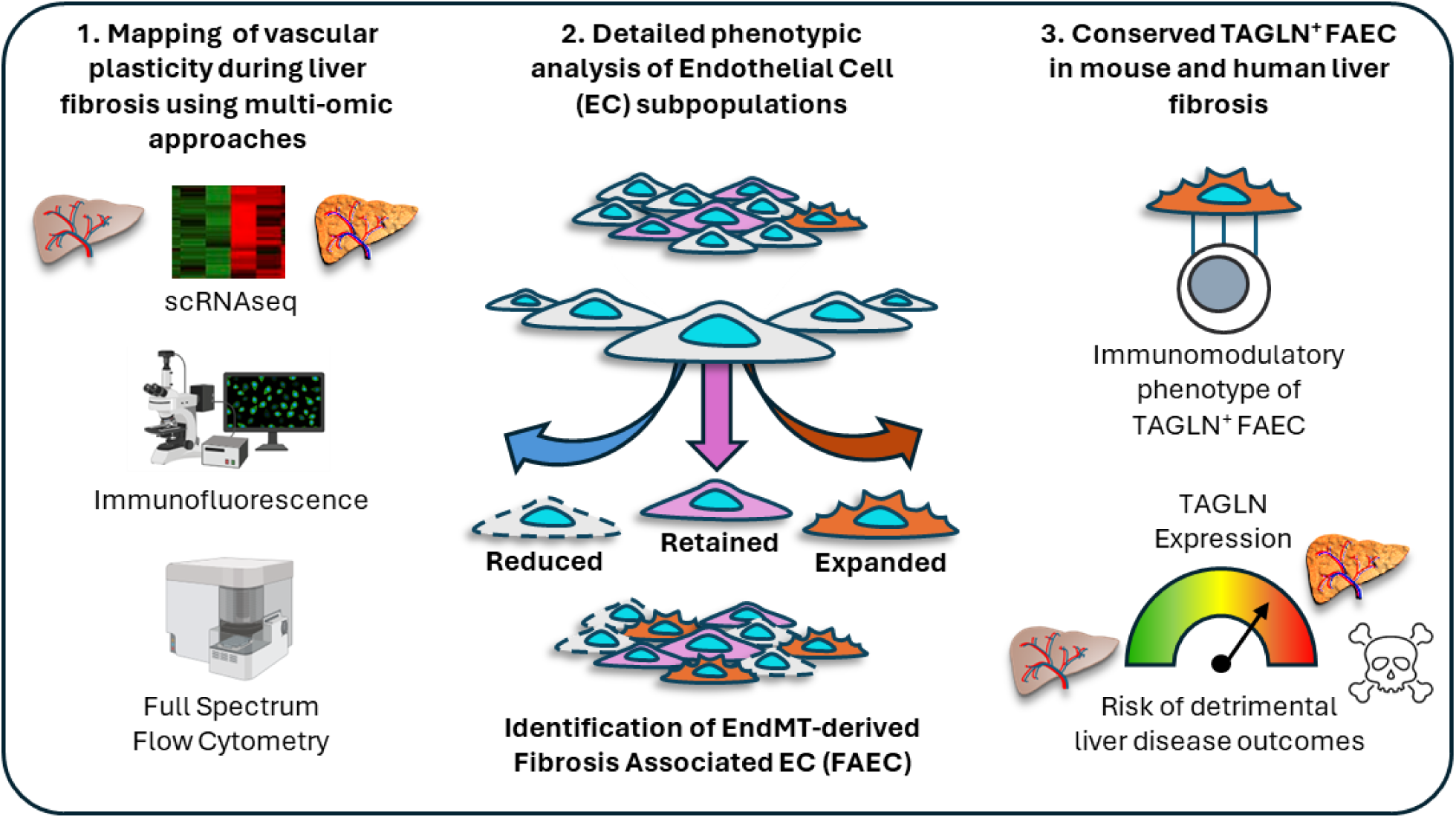

## INTRODUCTION

Fibrosis is the principal predictor of major adverse liver outcomes and of mortality in metabolic dysfunction-associated steatotic liver disease (MASLD). MASLD is the most common cause of chronic liver disease worldwide and its prevalence has risen in parallel with rising levels of obesity and type II diabetes(*1*). Despite major advances, including two drugs licensed in the United States(*2, 3*), the two main unmet clinical challenges in MASLD are sensitive and accurate diagnostics and treatments with clinically significant effect sizes, both of which rely on improved understanding of pathology including fibrogenesis.

Zonation of liver lobules containing specialised subpopulations of hepatic endothelial cells (EC), hepatocytes and immune cells, is critical for vascular, metabolic and immune homeostasis in the liver (*4–6*). Vascular zonation is dramatically impacted by chronic disease, with several studies documenting the changing EC landscape and the derivation of novel endothelial cell clusters between healthy and fibrotic liver tissue in mice (*7, 8*) and patients (*9, 10*). Endothelial-to-mesenchymal transition (EndMT) is a central endothelial plasticity pathway responsible for phenotypic vascular changes (*11–13*). It is defined as the process by which EC lose expression of endothelial lineage markers, such as CD31, and upregulate mesenchymal markers such as transgelin (TAGLN)(*14*), THY1.2(*15*) (also termed CD90.2) or platelet-derived growth factor receptor α (PDGFRα)(*16*). This process is predominantly driven through canonical transforming growth factor β (TGFβ) signalling and induces phenotypic changes such as enhanced expression of cytoskeletal and matrix-associated factors, including vimentin and collagen I and III (*8, 17*). Importantly, EndMT has been demonstrated to occur in both pre-clinical models of liver fibrosis (*11, 18*) and end-stage liver disease in patients (*11, 12*). The role of EndMT as a pathogenic driver, rather than a consequence of fibrogenesis, and its predictive value regarding disease progression remains ill-defined. We hypothesise that FAEC are EndMT-derived subpopulations that instigate both pro-fibrotic and pro-inflammatory pathways, driving the early transition from tissue injury to progressive liver fibrosis.

To address our hypothesis, we developed a multi-omic approach integrating single cell RNA sequencing (scRNAseq), immunofluorescent microscopy (IF) and full spectrum flow cytometry (FSFC) to identify and phenotype EC subpopulations during different stages of carbon tetrachloride (CCl_4_)- and Western diet (WD)-induced mouse models of liver fibrosis. We characterise the induction of two distinct EndMT-derived FAEC populations in our pre-clinical studies and translate our findings to three cohorts of MASLD patients. We show TAGLN^+^ FAEC are a common immunomodulatory EC subpopulation that share transcriptional characteristics, arising in both mouse models and patient samples, that are significantly elevated at early-stage disease prior to clinical appearance of fibrosis. Furthermore, the elevated expression *TAGLN* is a significant risk factor for liver decompensation events, even in the absence of fibrosis, and confers a significant additive effect for detrimental patient outcomes.

Taken together, we believe this study will pave the way for developing FAEC-specific diagnostic and therapeutic approaches to tackle progressive fibrotic liver disease.

## RESULTS

### Mapping EC plasticity in healthy and fibrotic liver tissue

We analyzed a publicly available dataset (GSE145086) enriched for hepatic immune cells, hepatic stellate cells (HSC) and EC in control C57BL/6 mice or following 2 and 4 weeks of CCl_4_-induced liver fibrosis(*19*) (Fig. 1A). We identified four hepatic EC clusters which expressed combinations of genes known to be characteristic for each zone (*4, 6*); Zone 1 EC (Cluster 8), Zone 2 EC (Clusters 0 and 1) and Zone 3 EC (Cluster 6; Fig. 1B). Each cluster also displayed distinct expression profiles for five EC-identity genes that are expressed as cell surface proteins: *Pecam1* (CD31), *Mcam* (MCAM), *Thbd* (TM), *Emcn* (EMCN) and *Lyve1* (LYVE1); (Fig. 1B-C). Immunofluorescence intensity profile for each of these markers, across the 200 µm distance between a portal and central vein confirmed this zonal expression in healthy liver tissue (Fig. 1D-E).

**Fig. 1.**
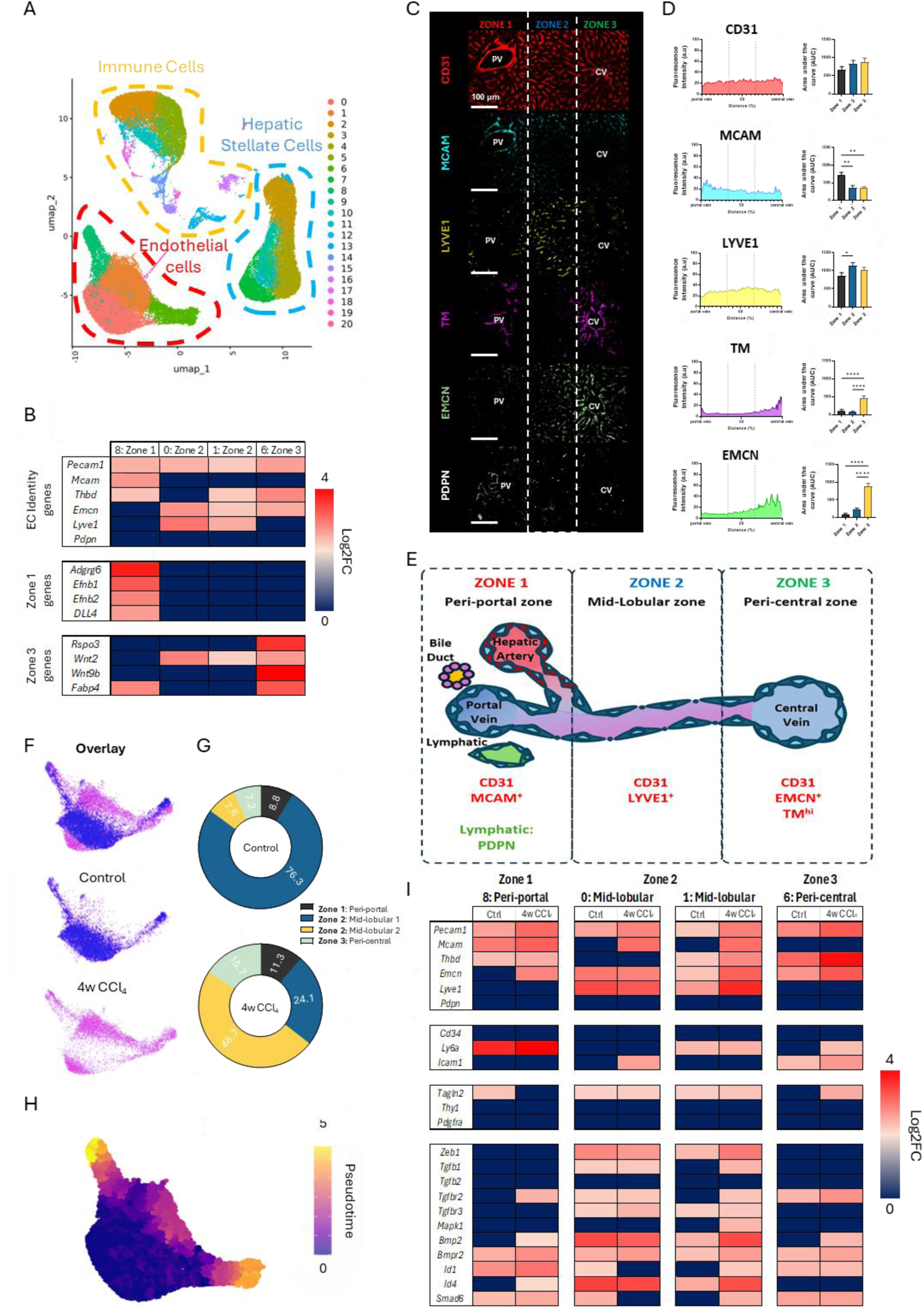
Characterisation of EC identity and plasticity markers in health and fibrogenesis. (A) UMAP of non-parenchymal cell scRNAseq data (GSE145086) for control (untreated), 2-and 4-week CCl_4_ treated male C57BL/6 mice(*19*). (B) Heatmap of gene expression for the four EC clusters (clusters 0, 1, 6 and 8) under baseline conditions identifying the zonation of EC identity markers (top panel) based on expression of established markers for peri-portal /Zone 1 (mid-panel) and peri-central/Zone 3 (bottom panel) EC. (C) Representative confocal images of healthy liver tissue stained for the EC identity markers CD31, melanoma cell adhesion molecule (MCAM), LYVE1, thrombomodulin (TM) and endomucin (EMCN). Podoplanin (PDPN) included as lymphatic marker. (D) Immunofluorescence intensity distribution across the portal-central vein axis and bar graphs of signal quantification for each zone for CD31, MCAM, LYVE1, TM and EMCN. Scale bars= 100μm. (E) Schematic summarising the zonal distribution of hepatic EC markers. (F) Visual overlay of EC clusters from scRNAseq for control mice (blue) and 4w CCl_4_ treatment (magenta). (G) Quantification of the abundance of each EC cluster in control (top panel) and 4w CCl_4_ treated mice (bottom panel). (H) Pseudotime analysis of progressive fibrogenesis from healthy, 2w and 4w CCl_4_-treated samples showing the transition away from cluster 0 mid-lobular EC toward clusters 1, 6 and 8. (I) Heatmap of gene expression for EC markers (top panels) and TGFβ-activated genes (bottom panels) for the four EC clusters comparing control and 4w CCl_4_ treatment groups.

After 4 weeks’ treatment with CCl_4_ to induce fibrosis, there was a marked shift in the abundance of mid-lobular EC populations during fibrogenesis (Fig. 1F and G), with pseudotime analysis showing reduction in the Zone 2 (Cluster 0) seen in health and expansion of what we have termed fibrosis-associated endothelial cells (FAEC) phenotype (Cluster 1) that transition towards Zone 1 and Zone 3 clusters (Fig. 1H). These expanding FAECs display enriched expression of a panel of EndMT-associated, TGFβ-driven, genes after 4 weeks’ CCl_4_ treatment (Fig. 1I). Notably, the expression of the EC identity genes also changes dynamically in response to CCl_4_ treatment (Fig. 1I), highlighting the plasticity of hepatic EC during fibrogenesis.

To study these shifts in EC identity and phenotype, we treated male C57BL/6 mice with peanut oil vehicle alone (sham), CCl_4_ for 4 weeks (4w; early fibrosis), 8 weeks (8w; established bridging fibrosis) or treated for an initial 4 weeks and then allowed to recover from tissue fibrosis for a further 4 weeks (Recovery Group; Fig. 2A-B).

**Fig. 2.**
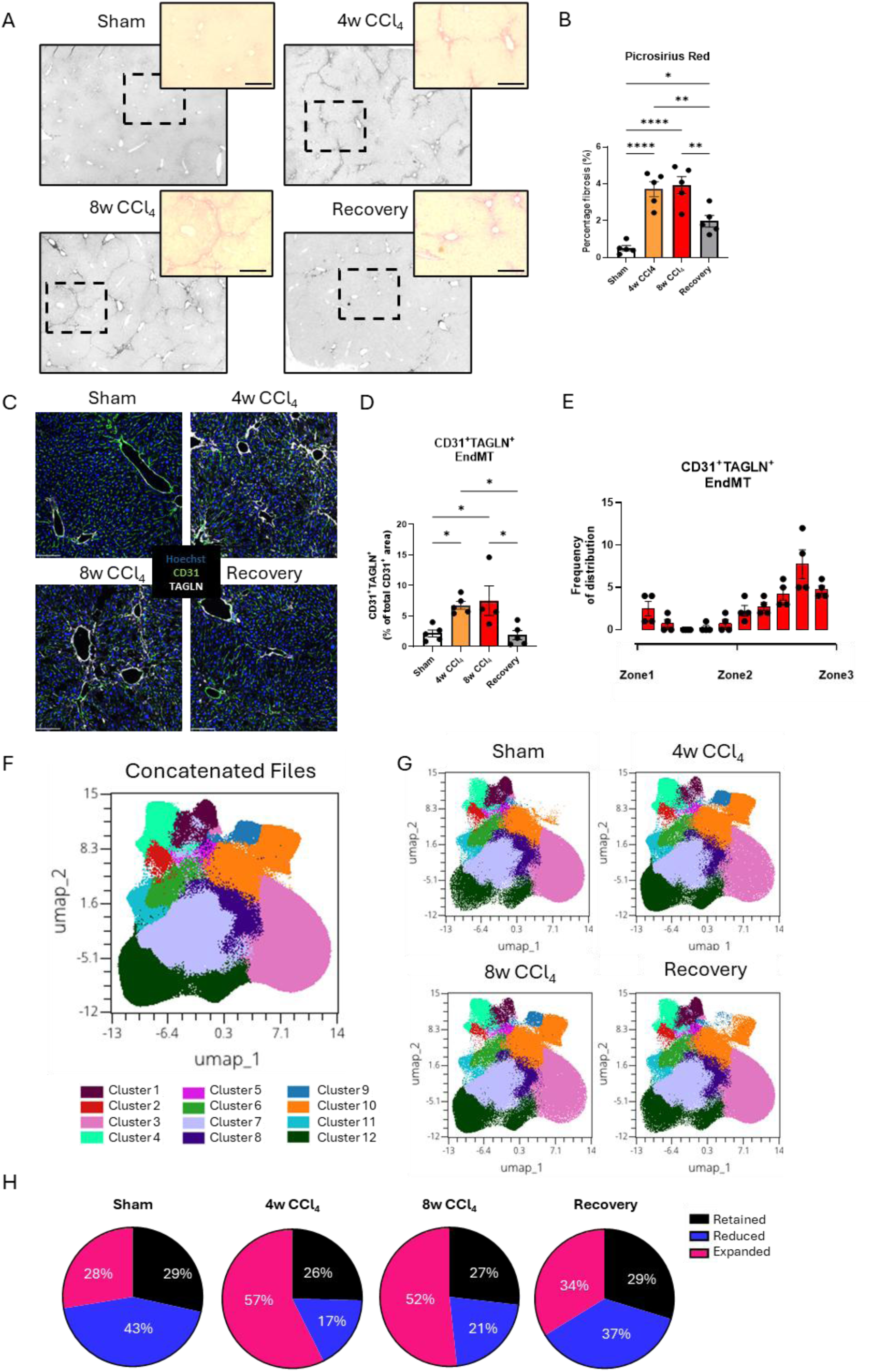
Mapping EC proteomic changes in response to CCl_4_-induced liver fibrosis and tissue recovery. (A) Representative images of picrosirius red staining of liver sections from Sham, 4- and 8-week CCl_4_ treated and recovery groups. Scale bar= 250μm. (B) Quantification of picrosirius red staining, expressed as a percentage (%) of total tissue area. (C) Representative confocal images of IF staining for Hoechst (blue), CD31 (green) and TAGLN (white) in Sham, CCl_4_-treated and recovery groups. Scale bar= 100μm. (D) Quantification of CD31^+^TAGLN^+^ EndMT cells expressed as a percentage of the total CD31^+^ area. (E) Bar graph of EndMT cell distribution across the portal vein to central vein axis. (F, G) UMAP of CD45^-^CD31^+^ cells from (F) control, CCl_4_-treated and recovery group samples displaying 12 distinct endothelial cell clusters and (G) CD45^-^CD31^+^ cells split by treatment group. (H) Pie charts of the percentage composition of total EC characterised as ‘Retained’ (black), ‘Reduced’ (blue) and ‘Expanded’ (pink) for each treatment group. For statistical analysis, one-way ANOVA tests with Tukey’s multiple comparisons test were used. N=5 animals/group. *; p<0.05, **; p<0.01, ****; p<0.0001.

Consistent with the transcriptional changes described above (Fig. 1), quantification of widefield IF images revealed significant changes in vascular activation showing the increase in co-localisation of vascular identity markers (CD31, TM and EMCN; Fig. 2C and Fig S1A-D) together with TAGLN (Fig. 2C & Fig S1A & E) resulting in a robust induction of CD31^+^TAGLN^+^ EndMT (Fig. 2D). EndMT and vascular activation were preferentially localized in Zone 3 peri-central regions (Fig. 2E). Taken together, we show that endothelial plasticity pathways are activated in distinct zones during liver fibrogenesis and tissue recovery, validating the suitability of this model to assess the role of EndMT in specific EC subpopulations leading to the derivation of FAEC phenotypes.

To observe phenotypic changes in multiple EC subpopulations simultaneously, a 14-marker panel was developed for FSFC(*20*) enabling us to profile the expression of 6 EC identity (CD31, MCAM, TM, EMCN, LYVE1 and PDPN), activation (CD34, ICAM1 and LY6A) and EndMT (TAGLN, THY1.2 and PDGFA) markers (Fig. S2A). All four treatment groups were concatenated and CD31^+^CD45^-^ ECs (Fig. S2B) were clustered based on expression of the identity markers, producing twelve populations (Concatenated data; Fig. 2F and split by treatment group Fig. 2G) that were subcategorized each based on change in abundance. **Reduced**: Significantly reduced abundance following CCl_4_ treatment compared to Sham, **Retained**: no significant change following CCl_4_ treatment compared to Sham and **Expanded**: significantly increased abundance compared to Sham (quantified in Fig. 2H). We show a striking alignment between the transcriptomic (Fig. 1G) and proteomic (Fig. 2H) plasticity data of hepatic EC but also highlight the considerable proteomic heterogeneity, which offers further insight into the phenotypic attributes of EC subpopulations during fibrogenesis and recovery.

### Phenotyping EC subpopulations during fibrogenesis and tissue recovery

‘Reduced’ ECs were the predominant subcategory of EC in healthy liver, comprising three clusters (1-3; Fig. 3A), which become less abundant following 4 and/or 8w CCl_4_ and returned towards basal levels in the Recovery group (Fig. 3B-D). All 3 Reduced EC clusters are LYVE1^+^ ICAM1^-^and can be distinguished based on expression of TM and EMCN (Fig. 3E), suggesting that the majority of these EC are situated in Zones 2 and 3 based on the expression profiles in Fig.1. These ECs had limited activation profiles, with LY6A induced in cluster 2 only (Fig. 3F), and no EndMT markers in response to CCl_4_ (Fig. S3A-C). The key phenotypic characteristics of the clusters in this category are summarised in Fig. 3G.

**Fig. 3.**
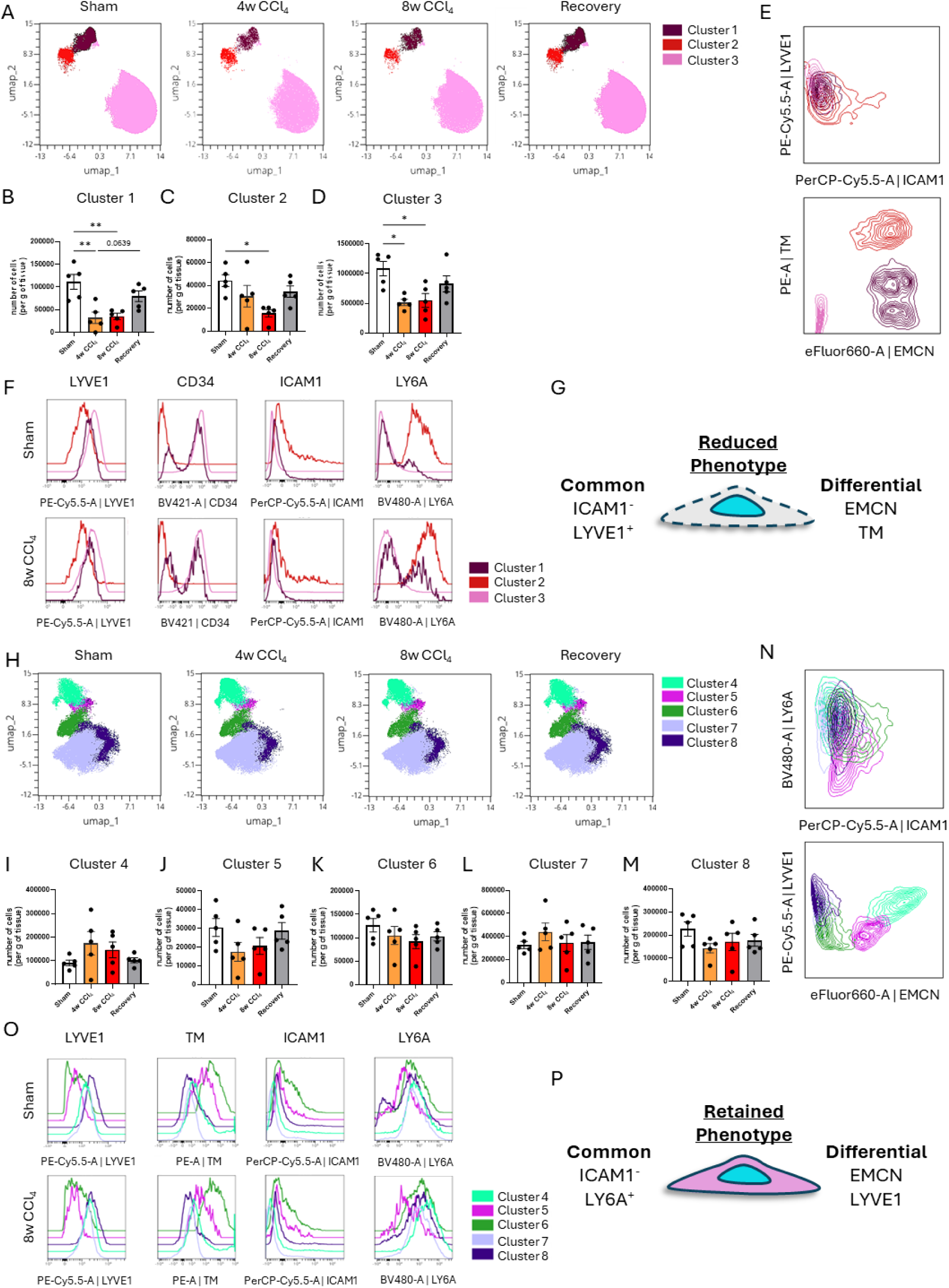
Phenotyping the characteristics of Reduced and Retained EC subpopulations by FSFC. (A) UMAP visualisation of three Reduced EC clusters; 1 (purple), 2 (red) and 3 (pink). (B-D) Quantification of the number of cells in each cluster expressed as number of cells per gram of liver tissue for (B) cluster 1, (C) cluster 2 and (D) cluster 3 per treatment group. (E) Contour plots of clusters 1, 2 and 3 for ICAM1 vs LYVE1 (top panel) and endomucin (EMCN) vs thrombomodulin (TM) (bottom panel). (F) Histograms of clusters 1, 2 and 3 comparing the median fluorescence intensity (MFI) of LYVE1, CD34, ICAM1 and LY6A in Sham (top panels) and 8w CCl_4_-treated animals (bottom panels). (G) Schematic summarising the common and differential traits of the three Reduced EC clusters. (H) UMAP visualisation of five Retained EC clusters; 4 (turquoise), 5 (light purple), 6 (green), 7 (violet) and 8 (dark purple). (I-M) Quantification of the number of cells in each cluster expressed as number of cells per gram of liver tissue for (I) cluster 4, (J) cluster 5, (K) cluster 6, (L) cluster 7 and (M) cluster 8 per treatment group. (N) Contour plots of clusters 4, 5, 6, 7 and 8 for ICAM1 vs LY6A (top panel) and EMCN vs LYVE1 (bottom panel). (F) Histograms of clusters 4, 5, 6, 7 and 8 comparing the MFI of LYVE1, TM, ICAM1 and LY6A in Sham (top panels) and 8w CCl_4_-tretaed animals (bottom panels). (G) Schematic summarising the common and differential traits of Retained EC clusters. For statistical analysis, one-way ANOVAs with Tukey’s multiple comparisons tests were used for figures B, D and I-M and a Kruskal-Wallis test with Dunn’s multiple comparisons test was used for figure C. N=5 animals/group. *; p<0.05, **; p<0.01.

Five ‘Retained’ clusters (4-8; Fig. 3H) did not change in abundance between healthy and CCl_4_-treated mice (Fig. 3I-M) although some clusters did express markers of activation and EndMT: MCAM^+^LY6A^+^TAGLN^+^ cluster 4 and PDGFRα^+^ cluster 6 (Fig. S3 D-H). Based on their expression profiles for Ly6A^+^ ICAM1^-^ and differential expression of Lyve1 and EMCN (Fig 3N-P) these EC do not appear to be associated to a specific zone.

Four Expanded EC clusters (9-12; Fig. 4A) significantly increased in abundance between healthy and CCl_4_-treated mice (Fig. 4B-E). We collectively define these clusters as FAEC as they are instigated in response to tissue fibrosis while returning to basal levels in the Recovery group. All expanded clusters expressed elevated markers of activation and EndMT (Fig. S3I-L). Notably, Clusters 9 and 10 expressed ‘classical’ EndMT markers, downregulating the expression of the EC identity markers EMCN and TM, respectively, while inducing the expression of THY1.2/PDGFRα, with increased inflammatory activation via ICAM1 (Fig. S3I&J). The most prevalent Expanded clusters, 11 and 12, displayed ‘non-classical’ EndMT phenotypes exhibiting both increased expression of EC identity markers, including CD31 and TM, together with enhanced TAGLN and LY6A expression (Fig. S3K&L). The idiosyncrasies of these 4 clusters are further emphasised by the differential contour plots (Fig. 4F) and histograms between the Sham and 8w CCl_4_ groups (Fig. 4G) and are summarised in schematics (Fig. 4H&I).

**Fig. 4.**
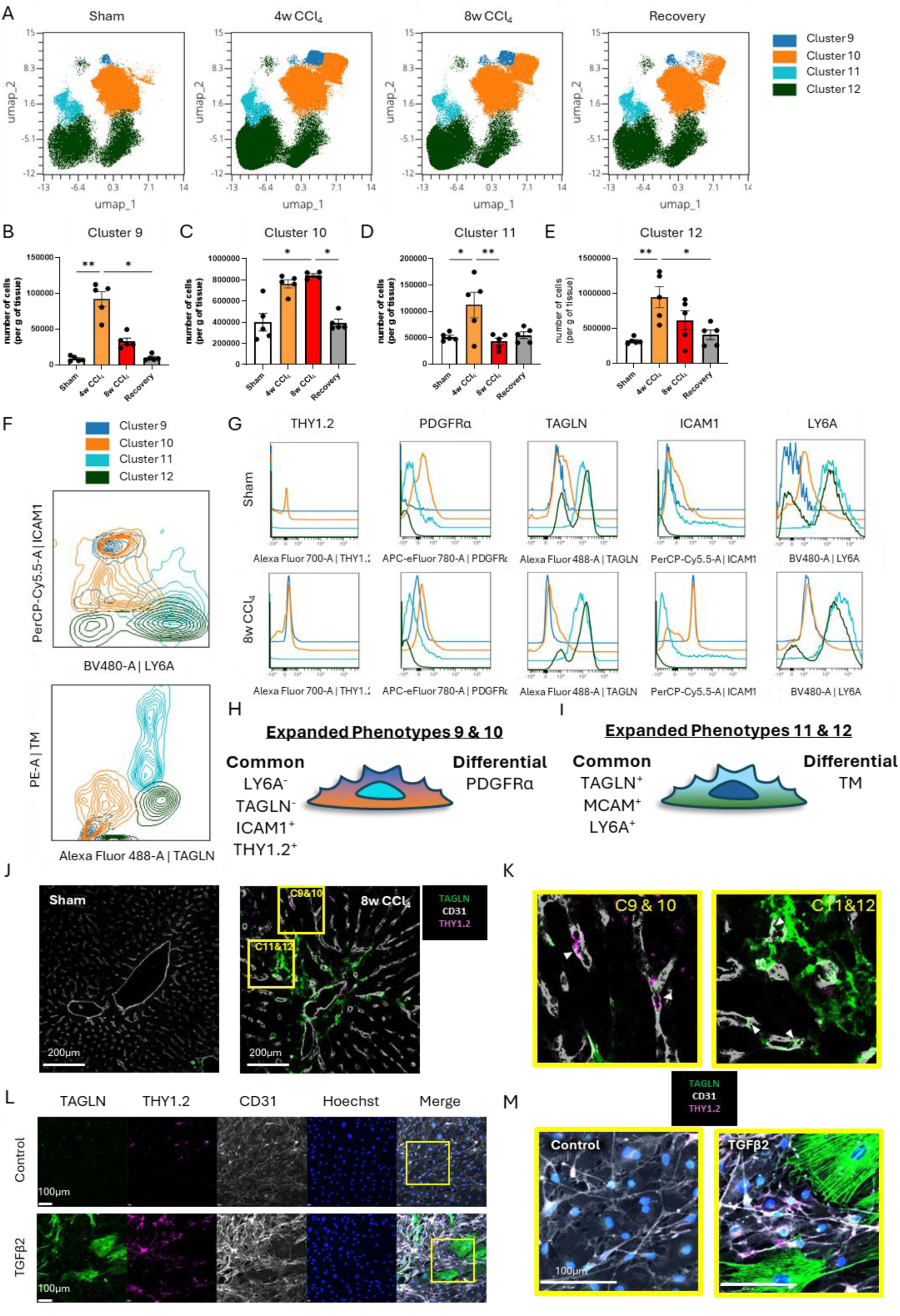
Phenotyping the characteristics of Expanded EC clusters by FSFC. (A) UMAP visualisation of four Expanded EC clusters; 9 (dark blue), 10 (orange), 11 (teal) and 12 (dark green). (B-E) Quantification of the number of cells in each cluster expressed as number of cells per gram of liver tissue for (B) cluster 9, (C) cluster 10, (D) cluster 11 and (E) cluster 12 per treatment group. (F) Contour plots of clusters 9, 10, 11 and 12 for LY6A vs ICAM1 (top panel) and transgelin (TAGLN) vs thrombomodulin (TM) (bottom panel). (G) Histograms of clusters 9, 10, 11 and 12 comparing the mean fluorescence intensity (MFI) for THY1.2, PDGFRα, TAGLN, ICAM1 and LY6A in Sham (top panels) and 8w CCl_4_-treated animals (bottom panels). (H, I) Schematics summarising the common and differential traits of (H) clusters 9 and 10 and (I) clusters 11 and 12. (J) Representative confocal images of IF staining for CD31 (white), THY1.2 (magenta) and TAGLN (green) in Sham and 8w CCl_4_-treated liver sections. Scale bar= 200μm (K) Magnified examples of CD31^+^THY1.2^+^ cells (left panel) and CD31^+^TAGLN^+^ cells (right panel). (L) Representative confocal images of IF staining for CD31 (white), THY1.2 (magenta), TAGLN (green) and Hoechst (blue) from isolated mouse EC cultured in the presence of either media alone or 10ng/ml TGFβ2 for 48h. Scale bar= 100μm. (M) Magnified examples of CD31^+^THY1.2^+^ cells and CD31^+^TAGLN^+^ cells in control (left panel) and TGFβ2-treated cells (right panel). Scale bar= 100μm. For statistical analysis, Kruskal-Wallis tests with Dunn’s multiple comparisons test were used for figures B and C and one-way ANOVA tests with Tukey’s multiple comparisons test were used for figures D and E. *; p<0.05, **; p<0.01.

In 8w CCl_4_- and Sham-treated mouse liver sections, THY1.2^+^CD31^+^ and TAGLN^+^CD31^+^ FAEC are mutually exclusive cells within Zone 2 and Zone 3 regions of the liver, respectively (Fig. 4J&K). To determine whether TGFβ-induced EndMT can give rise to both FAEC populations, isolated healthy mouse liver EC stimulated *in vitro* with 10 ng/ml TGFβ2 for 48 hours also showed differential expression of THY1.2 and TAGLN (Fig. 4L&M).

Together, these data demonstrate EC heterogeneity and plasticity during fibrosis and tissue recovery and identify two distinct types of FAEC. While THY1.2^+^ and TAGLN^+^ FAEC may arise from a common TGFβ-driven EndMT pathway, they may also originate from separate healthy EC subpopulations. To test this assertion, we addressed the potential for Reduced clusters to transition towards Expanded FAEC clusters using Wishbone trajectory analysis.

### Derivation of FAEC using Wishbone trajectory mapping

To determine whether EC in Reduced clusters transition towards Expanded FAEC, we first assessed their overlapping traits. Reduced clusters 1 and 3 and Expanded clusters 9 and 10 overlapped in expression of LYVE1 (Fig. 5A) and LY6A (Fig. 5B) in Sham animals and displayed marked plasticity following treatment. Notably, a subset of cluster 10 became THY1.2^+^ (Fig. 5B) while there was a complete transition for cluster 9 towards a THY1.2^+^ phenotype in response to CCl_4_ treatment (Fig. S3I&J). Wishbone trajectory analysis for the four clusters at all fibrosis stages, showed that LYVE1^high^ ICAM1^low^ THY1.2^low^ EC converge at a branch point which enables them to either shift towards a FAEC phenotype of clusters 9 and 10 in response to 8w CCl_4_ treatment or return to a healthy phenotype during recovery (Fig. 5C). The inverse relationship of these markers is highlighted in the trajectory heatmaps showing that LYVE1 and THY1.2 appear to be mutually exclusive, while ICAM1 expression is enhanced at a later stage of transition to FAEC (Fig. 5D). The putative transition pathway is summarised schematically (Fig. 5E).

**Fig. 5.**
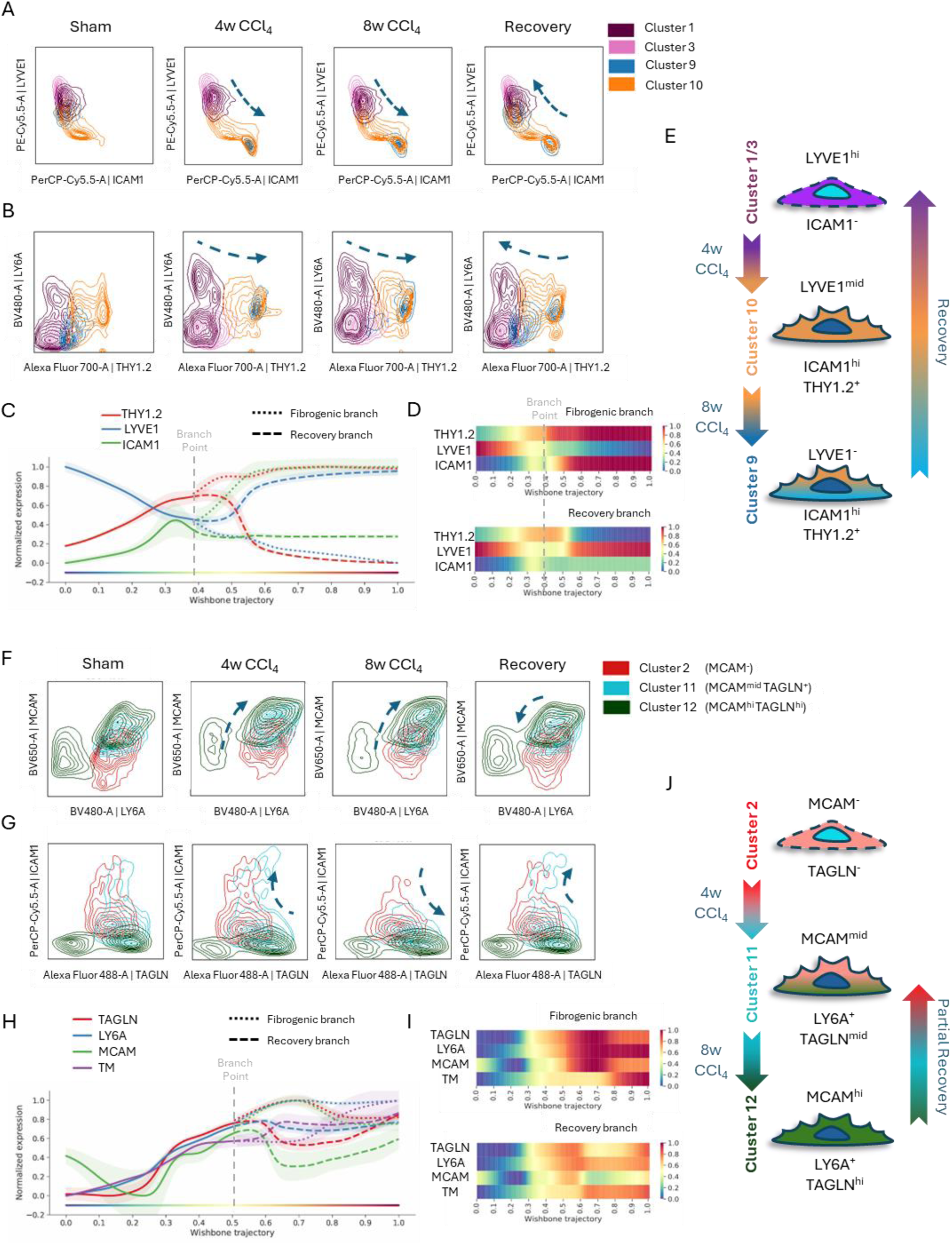
Derivation of Expanded FAEC from Reduced EC clusters through acquisition of EndMT-traits. (A, B) Contour plots of clusters 1, 3, 9 and 10 for (A) ICAM1 vs LYVE1 and (B) THY1.2 vs LY6A. Arrow denotes the shifting phenotypic profile of the clusters for the different treatment groups. (C) Wishbone trajectory analysis of clusters 1, 3, 9 and 10 for THY1.2 (red), LYVE1 (blue) and ICAM1 (green) (D) Heatmap displaying the expression profile of the Fibrogenic branch (top panel) or Recovery branch (bottom panel) for THY1.2, LYVE1 and ICAM1. (E) Schematic summarising the trajectory and profile of clusters 1, 3, 9 and 10 during fibrogenesis and tissue recovery. (F, G) Contour plots of clusters 2, 11 and 12 for (F) LY6A vs MCAM and (G) transgelin (TAGLN) vs ICAM1. Arrow denotes the shifting phenotypic profile of the clusters for the different treatment groups. (H) Wishbone trajectory analysis of clusters 2, 11 and 12 for TAGLN (red), LY6A (blue), MCAM (green) and TM (purple). (I) Heatmap displaying the expression profile of the Fibrogenic branch (top panel) or Recovery branch (bottom panel) for TAGLN, LY6A, MCAM and thrombomodulin (TM). (J) Schematic summarising the trajectory and profile of clusters 2, 11 and 12 during fibrogenesis and tissue recovery.

Similarly, Reduced Cluster 2 and the two TAGLN^+^ FAEC clusters 11 and 12 displayed overlapping expression of MCAM and LY6A (Fig. 5F) and TAGLN (Fig. 5G), which were all elevated following 4 and 8 weeks’ CCl_4_ treatment. Incremental acquisition of the TAGLN^+^MCAM^+^ FAEC phenotype during fibrosis is only partially restored after recovery, suggesting that these FAEC may persist in the tissue even after the insult is removed (Fig. 5H). The trajectory heatmap highlights the persistence of TAGLN, LY6A and MCAM in both the fibrogenic and recovery branches (Fig. 5I), suggesting these FAEC have decreased plasticity, impairing their capacity to recover a healthy phenotype (Fig. 5J).

### Immunomodulatory function of FAEC

To assess the influence of these two distinct FAEC subpopulations on the inflammatory response within the liver during fibrogenesis and recovery, we explored the transcriptomic signatures of EC enriched for either *Thy1* or *Tagln2* in the scRNAseq dataset and compared them with the phenotypic characteristics in our FSFC data. Abundance of these cell types was similar: *Thy1*^+^ FAEC comprising ∼1% of total EC by scRNAseq and THY1.2^+^ FAEC ∼5% of total EC by FSFC and, *Tagln^+^* (scRNAseq) and TAGLN^+^ (FSFC) FAEC comprising ∼20%. Gene expression in *Thy1^+^* FAEC was indicative of leukocyte interaction and activation (Fig. 6A) and THY1.2^+^ FAEC expressed increased levels of ICAM1 following CCl_4_ treatment in both cluster 9 and 10, and reduced levels of the anti-inflammatory glycoprotein EMCN in cluster 9 only (Fig. 6B). *Tagln^+^* FAEC displayed more specific enrichment for metabolic and inflammatory genes (Fig. 6C) and TAGLN^+^ FAEC were characterised by upregulation of MCAM and LY6A (Fig. 6D). In line with the localisation and phenotype of FAEC CD45^+^ leukocytes (Fig. 6E) are preferentially recruited to peri-central regions (Fig. 6F) following 4 and 8 weeks’ CCl_4_ treatment.

**Fig. 6.**
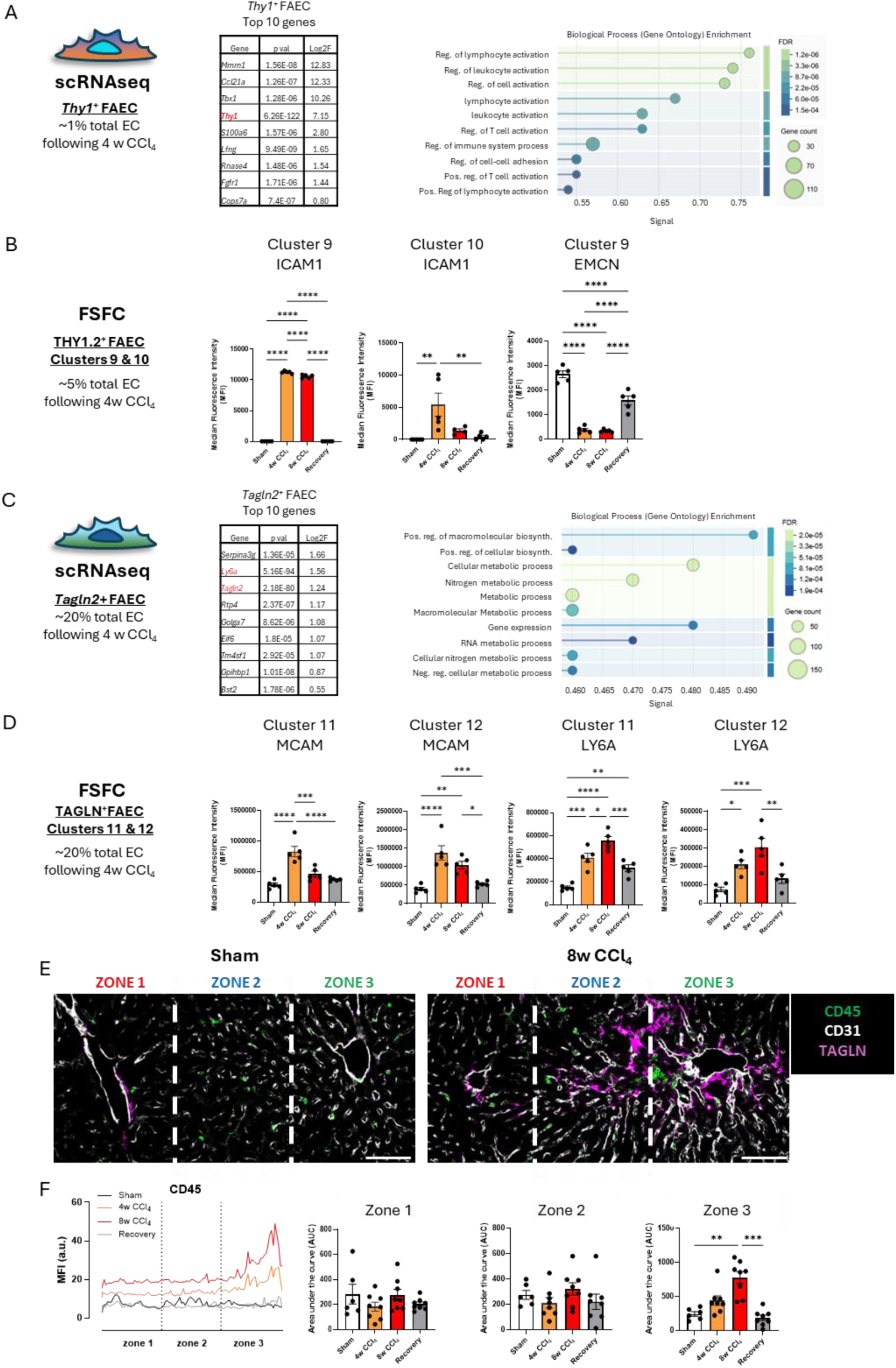
Transcriptomic and proteomic phenotype of THY1^+^ and TAGLN^+^ FAEC. (A) ScRNAseq analysis of *Thy1* expressing liver EC from 4w CCl_4_-treated mice compared to all other EC. Table of the top 10 enriched genes and gene ontology analysis of putative biological processes enriched in *Thy1*^+^ EC based on their differentially expressed genes. (B) FSFC analysis of median fluorescence intensity (MFI) of Thy1.2^+^ FAEC for ICAM1 and EMCN in cluster 9 and cluster 10. (C) ScRNAseq analysis of *Tagln2* expressing liver EC from 4w CCl_4_-treated mice compared to all other EC. Table of the top 10 enriched genes and gene ontology analysis of putative biological processes enriched in *Tagln2^+^* EC based on their differentially expressed genes. (D) FSFC analysis of median fluorescence intensity (MFI) of TAGLN^+^ FAEC for MCAM and LY6A in cluster 11 and 12. (D) Representative confocal images of IF staining for CD45 (green), CD31 (white) and TAGLN (magenta) in Sham and 8w CCl_4_-treated livers. Scale bar= 100μm. (E) Profile plot of CD45^+^ MFI across the portal-central vein axis in Sham, CCl_4_-treated and recovery experimental groups. (F) Quantification of the profile plot area under the curve for Zone 1, Zone 2 and Zone 3. (a.u.: arbitrary units). For statistical analysis, one-way ANOVA tests with one-way ANOVA tests with Tukey’s multiple comparisons tests were used. N=5 animals/group. *; p<0.05, **; p<0.01, ***; p<0.001, ****; p<0.0001.

### TAGLN^+^ FAEC are conserved in Western diet-fed mice and MASLD patients

Having characterised the induction of FAEC subpopulations in a murine CCl_4_-induced liver fibrosis, we sought to determine if FAEC were also an early characteristic of a murine dietary model of MASLD induced by feeding animals a Western Diet (WD). There was significant uptake of lipid within the liver (Fig. 7A and B) and moderate hepatic fibrosis (Fig. 7C and D). In keeping with the instigation of mild tissue fibrosis, compared to the CCl_4_ model, the impact on vascular architecture (Fig. S4A and B) and EC identity markers was less pronounced, but still resulted in significantly increased EMCN, TM and TAGLN following 12w of WD (Fig. S4C).

**Fig. 7.**
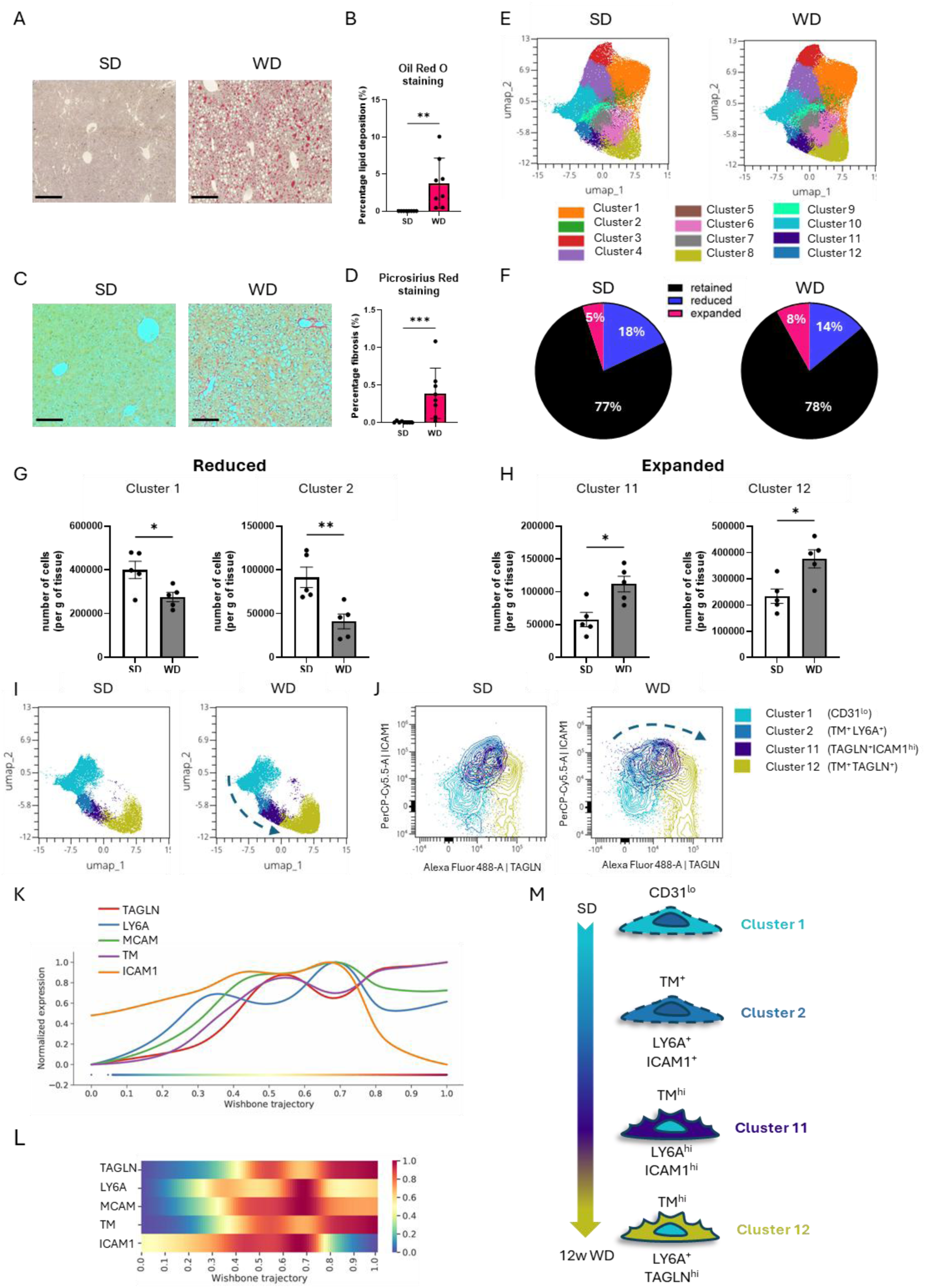
Expansion of TAGLN^+^ FAEC in a 12w Western Diet model of MASLD. (A) Representative images of Oil Red O staining of liver samples from 12w standard diet (SD) and Western diet (WD) fed mice. Scale bars = 250 μm. (B) Quantification of Oil Red O staining expressed as a percentage (%) of lipids detected over the total tissue area. (C) Representative images of Picrosirius Red staining of liver samples from 12w SD and WD fed mice. Scale bars = 250 μm. (D) Quantification of Picrosirius Red staining expressed as a percentage (%) of collagen detected over the total tissue area. (E) UMAP plots of CD45^-^CD31^+^ EC clusters from SD or WD fed mice. (F) Pie charts of the percentage composition of total EC characterised as ‘Retained’ (black), ‘Reduced’ (blue) and ‘Expanded’ (pink) for each diet group. (G, H) Quantification of the number of cells per gram of liver tissue for (G) Reduced cluster 1 and cluster 2 and (H) Expanded cluster 11 and cluster 12. (I) UMAP of cluster 1 (turquoise), cluster 2 (blue), cluster 11 (purple) and cluster 12 (yellow) from SD or WD groups. (J) Contour plots of transgelin (TAGLN) vs ICAM1 for clusters 1, 2, 11 and 12 for SD or WD groups. Arrow denotes the shifting phenotypic profile of the clusters from SD to WD. (K) Wishbone trajectory analysis of clusters 1, 2, 11 and 12 for TAGLN (red), LY6A (blue), MCAM (green), TM (purple) and ICAM1 (orange). (L) Heatmap of the expression profile of each marker. (M) Schematic summary of the trajectory and profile of clusters 1, 2, 11 and 12. For statistical analysis, unpaired t-tests were used for figures B, G, H and a Mann-Whitney test was used for figure D. N=5-8 animals/group. *; p<0.05, **; p<0.01, ***; p<0.001.

Assessment of EC subpopulations by FSFC distinguished twelve hepatic EC clusters (Fig. 7E). Consistent with our earlier findings (Fig. 2F), we identified phenotypically distinct Reduced, Retained and Expanded subpopulations (Fig. S5A-L) albeit the differences in proportion were more subtle than CCl_4_ (Fig. 7F). Reduced clusters 1 and 2 (Fig. 7G), and Expanded clusters 11 and 12 (Fig. 7H) shared overlapping characteristics when compared on the UMAP (Fig. 7I), such as ICAM1 and TAGLN expression (Fig. 7J). Wishbone analysis revealed a shared trajectory towards a TAGLN^+^ FAEC phenotype (Fig. 7K and L), which occurred more gradually than in response to CCl_4_ (Fig. 7M).

Finally, we examined whether the induction of TAGLN expression was associated with disease severity and patient outcomes in human MASLD. In liver biopsy samples from patients who underwent bariatric surgery, there was a significant expansion of both the number of *TAGLN*^+^ EC and expression of *TAGLN*^+^ in patients with compared to those without fibrosis (Fig. S6A). IF staining for TAGLN^+^CD31^+^ EndMT cells (Fig. S6B) showed a significant increase in patients with early stages (F0/1) of disease (Fig. S6C) compared to healthy control. In keeping with the phenotype of TAGLN^+^ FAEC in murine models, we found enrichment of *MCAM*, *TAGLN* and *THY1* (Fig. S6D) with GO analysis showing similar enrichment to the mouse data in vascular remodelling, pro-fibrotic and pro-inflammatory pathways (Fig. S6E). In spatial transcriptomic analysis of four patient biopsies (Supplementary Table 1). TAGLN^+^ FAEC co-localised with ECM and inflammatory macrophages with regions of fibrosis (Fig. 8A). In line with patient scRNAseq data (Fig. S7A-C), the appearance of TAGLN^+^ FAEC was apparent at early-stage disease before observable fibrosis and persisted in patients with cirrhosis (Fig. S7A). Comparison of TAGLN^+^EC vs TAGLN^-^ EC transcriptomic signatures revealed that pro-fibrotic (Collagen) and pro-inflammatory (ICAM) pathways were preferentially enriched in TAGLN^+^ FAEC (Fig. 8B and Fig. S7B). Detailed CellChat analysis identified the expression of *THBS, LAMA* and *COL* as potentially unique sender/receiver partners (Fig. 8C and Fig. S7C).

**Fig. 8.**
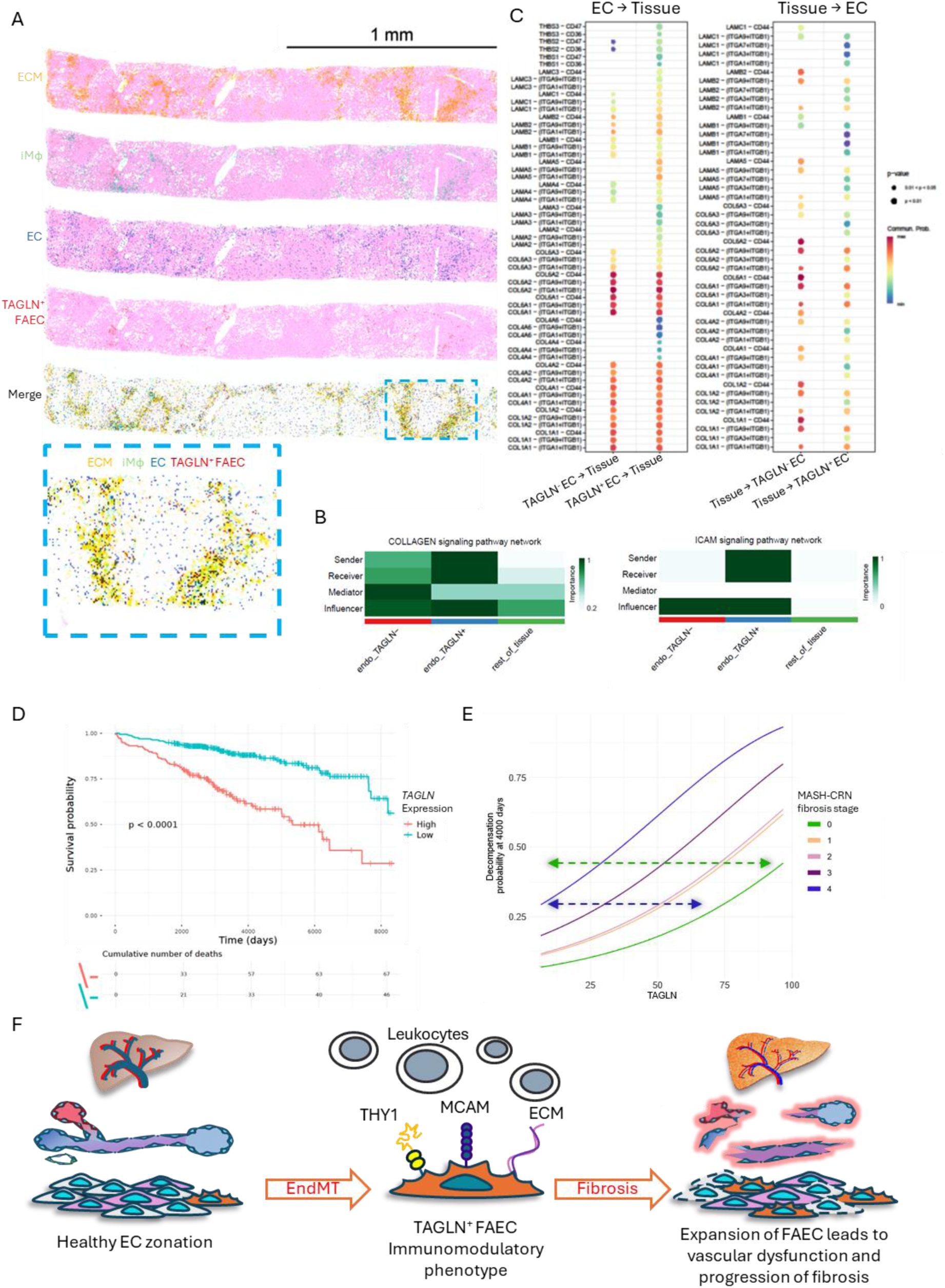
Expression of TAGLN^+^ FAEC is a conserved feature of MASLD with the potential to be a biomarker of progressive disease. (A) Visualization of spatial transcriptomic clusters for *k*-means shows enrichment data for a patient with Ishak score of 2-3 identifying extracellular matrix (ECM; yellow), inflammatory macrophages (iMϕ), all endothelial cells (EC; blue) and TAGLN^+^EC (red). Zoomed in region highlighted in blue box (B) Comparison between TAGLN^-^ EC and TAGLN^+^EC for collagen and ICAM signalling network displayed as heatmaps. (C) CellChat analysis comparing enriched of putative interaction patterns from EC to tissue and tissue to EC. (D) Kaplan–Meier survival curves comparing patients stratified into high and low TAGLN expression groups (p-value of log-rank test comparing groups) in the biopsy subset of the SteatoSITE cohort (n=505). (E) Cumulative incidence functions for liver-related events at 4000 days post-biopsy as a function of TAGLN expression, stratified by pathologist-scored MASH-CRN fibrosis stage (n=435; F0 (green), F1 (orange), F2 (pink), F3 (purple) and F4 (blue). Scale bar = 50 μm. For statistical analysis a one-way ANOVA with Dunnett’s multiple comparisons test was used. **; p<0.01. (F) Summary schematic of the role of FAEC in the development and progression of vascular dysfunction, fibrosis and MASLD.

To evaluate the potential clinical impact of elevated TAGLN expression in MASLD patients, we utilised the SteatoSITE(*21*) database to assess patient outcomes. High *TAGLN* expression (above 26.76 cpm) in liver biopsies revealed patients at a significantly elevated risk of all-cause mortality compared to those with lower expression levels (Fig. 8D). Furthermore, patients with high levels of *TAGLN* but no fibrosis (Fig. 8E, F0; green line) had a significantly higher risk of hepatic decompensation than patients with cirrhosis but low *TAGLN* expression (Fig. 8E, F4; blue line), suggesting that *TAGLN* expression has the potential to be an early and fibrosis-independent risk factor of detrimental patient outcomes.

Overall, our data elucidate the derivation of two distinct FAEC subpopulations by integrating scRNAseq and IF with a novel FSFC analysis of EC plasticity. We show that TAGLN^+^MCAM^+^ EC are the predominant FAEC phenotype in two mechanistically distinct models of murine liver fibrosis and are also present in patients with MASLD. Notably, these FAEC share common pro-fibrotic and pro-inflammatory phenotypes, which have potential immunomodulatory functions by co-localising with leukocytes (summarized in Fig. 8F). Finally, we reveal that the induction of *TAGLN* expression is significantly associated with detrimental clinical outcomes in liver disease patients, independent of fibrosis severity, supporting the hypothesis that TAGLN^+^ FAEC play an underappreciated role in disease pathogenesis.

## DISCUSSION

The role of vascular heterogeneity in health and disease has come into sharp focus in recent years, with considerable advances in single-cell transcriptomic(*7, 22*) and proteomic(*6*) profiling revealing remarkable insight into organotypic and pathological endothelial heterogeneity. In this study, three broad categories of liver EC in liver fibrosis: ‘Reduced’, ‘Retained’ and ‘Expanded’ have been defined and phenotyped. We find that the Reduced clusters that express CD31^+^LYVE1^+^MCAM^neg^ (some also expressed EMCN) localise to mid-lobular (Zone 2) and peri-central (Zone 3) regions, the same regions where the Expanded cells arise from, in keeping with known CCl_4_ damage pattern(*23, 24*). Using Wishbone trajectory analysis, we were able to show that they can change their phenotype and contribute to the development of Expanded populations. All Expanded clusters expressed EndMT traits as well as enhanced inflammatory profiles in response to CCl_4_-induced liver fibrosis. These EC clusters were the most highly plastic of all the EC subpopulations assessed, dramatically changing their abundance and phenotype in response to both CCl_4_ timepoints and following tissue recovery, thus we designated them as FAEC. Finally, the association of these markers with diseases severity and outcome in human disease strongly suggest translational potential of these findings(*25*).

EC are highly plastic(*26*) and their regulation of reparative or pathogenic pathways (*27, 28*) represent new diagnostic and therapeutic potential in human disease. We and others have determined that TGFβ-induced EndMT cells are an important component of the fibrogenic hepatic vascular niche(*11, 12, 29, 30*), however, the functional consequences of these phenotypic changes remain elusive. There is currently no consensus on how to define EC heterogeneity in the liver, and the application of multiple techniques has led to a variety of distinct subpopulations, ranging from immunohistochemistry studies(*31*) detailing type 1 and type 2 LSEC to single-cell RNAseq identifying five to seven endothelial populations in mice(*32*) and humans(*10*). A key advancement of our study is the complementary use of available scRNAseq data with multiple identity marker and high-throughout proteomic phenotyping through FSFC. While previous studies applied lineage tracing approaches to validate the occurrence of EndMT(*7, 12, 17, 33*) in several tissues and disease states, however, they are restricted to an individual identity marker. In contrast, our approach provides a more comprehensive analysis of EC plasticity, giving large-volume, single-cell data with multiplexing capabilities, high-dimensional reduction analysis and predictive algorithms, such as Wishbone trajectory(*20, 34*), to trace the derivation of novel FAEC populations.

The most prominent distinguishing feature of these FAEC clusters was the expression of either THY1.2^+^ or TAGLN^+^. THY1.2^+^ FAEC express higher levels of ICAM1 and do not express EMCN and LYVE1. THY1.2 is a member of the immunoglobulin supergene family and is commonly associated with smooth muscle cells. EC expressing *Thy1* are associated with inflammation-induced angiogenesis(*35*) and trans-endothelial leukocyte migration(*36*). ICAM1 is a key regulator of leukocyte rolling and adhesion to LSEC(*37*). Conversely, loss of EMCN and LYVE1 can also influence immune cell interactions, as both are documented to play protective roles in preventing aberrant leukocyte recruitment, respectively (*38, 39*).

TAGLN^+^ FAEC were not only the most abundant but were also conserved across species and displayed reduced plasticity, which may account for their persistence across all stages of liver disease assessed. TAGLN, a member of the calponin superfamily, is an actin-binding protein which can regulate the cytoskeleton(*40*) and ectopic TAGLN expression is hallmark of EndMT (*26, 41*). The upregulation of MCAM may point towards its role as an adhesion molecule that promotes cell-ECM and cell-cell interactions(*42*). MCAM is a well-characterised biomarker for advanced liver disease(*43*) and it is known to induce pro-coagulant responses in metastatic tumours(*44*). Upregulation of Ly6A is consistent with previous reports detailing the expansion of Ly6A^+^ LSEC during liver injury (*45, 46*). Notably, TAGLN^+^ MCAM^hi^ FAEC have a conserved phenotype that is induced in early stages of liver injury in both CCl_4_- and 12-week WD-induced mouse models and in patients without observable fibrosis, suggesting that the appearance of TAGLN^+^FAEC could be a novel, sensitive marker of disease progression.

TAGLN^+^ FAEC co-localise with regions enriched for ECM and infiltrating leukocytes in both mouse and human fibrosis. In the large SteatoSite cohort of histologically confirmed MASLD (*21*), we find that high hepatic *TAGLN* expression is significantly associated with adverse clinical outcomes, at all disease stages extending prognostic insight beyond fibrosis severity.

Taken together, we have identified distinct EndMT-derived FAEC with defined localisation and characteristics which have the capacity to reshape the vascular and immune landscape of the liver. TAGLN^+^ FAEC represent an exciting opportunity to develop new diagnostic and therapeutic approaches to tackle the growing clinical burden of chronic liver disease.

## MATERIALS AND METHODS

### Animals

All procedures used in this study were approved by the Animal Use and Care Committee of Queen Mary University of London and were in accordance with national and international regulations (Project license: PP0505769).

8-10 weeks old wild-type (WT) C57BL/6 male mice were purchased from Charles River Laboratories UK Ltd. (Kent, UK). For all *in vivo* experiments, power calculations were performed using preliminary data to calculate the appropriate number of animals to be included in the study.

### Murine models of liver injury

Carbon tetrachloride (CCl_4_; 289116-100ML, Sigma Aldrich) was diluted 1 in 5 in peanut oil (P2144-250ML, Sigma) and was administered with an intraperitoneal (IP) injection at a dose of 0.8mg/Kg in 100μl. Injections were administered twice a week for 2, 4 or 8 weeks. For the recovery experiments, mice were injected with CCl_4_ for 4 weeks and were afterwards left to recover for an additional 4 weeks. As a control, no treatment or Sham injections of 100μl peanut oil were used. To model a western diet (WD) and induce hepatic steatosis and fibrosis, mice were fed the modified Clinton-Cybulsky cholesterol diet (58R6, IPS Product Supplies) with the addition of carbohydrates (42g/L) consisting of 55% fructose (F0127-1KG, Sigma-Aldrich) and 45% sucrose (S9378-1KG, Sigma-Aldrich) in the drinking water for 12 weeks At each stated time point, animals were sacrificed, followed by cardiac perfusion with 3ml sterile deionized phosphate-buffered saline (PBS) (BR0014G, ThermoFisher Scientific) with 0.5% bovine serum albumin (BSA) (BP9702-100, Fisher Scientific) and 2mM EDTA (E7889-100ML, Sigma Aldrich) through the LV. Individual liver lobes were removed for immunohistochemistry, tissue digestion for tissue culture or flow cytometry and RNA extraction.

### Immunohistochemistry

Tissues were fixed for 2 hours in 4% paraformaldehyde before being prepared for either OCT or paraffin embedding. Tissue sections were cut at 10μm or 5μm, respectively. Samples were initially blocked in 3% BSA diluted in PBS +0.1% Tween 20 (P1379-100ML, Sigma) (PBST) for 1 hour at room temperature (RT). The samples were incubated with primary antibodies (Supplementary table 2) overnight (diluted in 1% BSA). They were then washed 3 times with PBST and incubated with appropriate secondary antibodies (Supplementary table 2) for 1 hour at RT. The slides were washed and mounted with FluoroMount G (00-4958-02, Invitrogen). Images were acquired using an inverted confocal laser scanning microscope at a 10x magnification.

### Picrosirius Red Staining

Paraffin-embedded sections were deparaffinised with Histoclear solution (NAT1330, Scientific Laboratory Supplies) and rehydrated. For staining, the Abcam Picrosirius red stain kit was used (ab245887, Abcam). The slides were then incubated in phosphomolybdic acid for 8 minutes and were transferred to picrosirius red stain for 60 minutes. They were then washed 3 times with acetic acid and were mounted using the Entellan (107960, Merck) mounting medium. The samples were visualised using the Olympus CKX53 light microscope (Olympus, Japan) and images were acquired at 4x magnification.

To analyse the data, the whole region from 2 sections per sample was quantified. The percentage (%) of picrosirius red-positive area was calculated using Image J and the colour deconvolution macro(*47*).

### Tissue Digestion

Liver samples were mechanically disrupted until homogenised using scissors. The homogenates were then further digested enzymatically in 3ml digestion buffer containing Hank’s Balanced Salt Solution (HBSS) with CaCl_2_ and MgCl_2_ (14025092, Gibco) with 20mM N-2-hydroxyethylpiperazine-N’-2-ethanesulfonic acid (HEPES) (15630-080, Gibco), 2mg/ml collagenase (17018-029, Gibco), 8U/ml dispase II (04942078001, Roche) and 50U/ml DNase (18047-019, Invitrogen). For digestion, the homogenates were incubated at 37°C for 30 minutes on a shaker. The samples were then filtered twice through a 70μm cell sieve (431751, Scientific Laboratory Supplies). The cell pellets were then resuspended for tissue culture or full spectrum flow cytometry.

### Staining for Full Spectrum Flow Cytometry (FSFC)

Single cell suspensions were transferred to a V-shaped 96-well plate (50μl/ well). The cells were washed with 200μl PBS and were resuspended in 50μl Fc blocking solution (130-092-575, Miltenyi Biotec) containing the Live/Dead^TM^ Fixable Blue stain (L23105, Invitrogen) at a dilution of 1 in 5,000 for 30 min at 4°C. The cells were washed twice with 200μl PBS and were then incubated with cell surface binding antibodies for 30 min at 4°C. FSFC primary antibody details are listed for the endothelial cell panel (Supplementary table 3). The samples were then washed with PBS and centrifuged at 400g for 2 minutes at 4°C. For fixation and permeabilization, the cell pellets were resuspended in the intracellular fixation and permeabilization buffer (00-5521-00, Invitrogen) and the plate was incubated for 20 min at room temperature (RT). The samples were then washed with 200μl permeabilization wash buffer (00-8333-56, Invitrogen) and centrifuged at 400g for 2 minutes. The cell pellets were resuspended in the intracellular binding antibody solution and incubated for 30 minutes at RT. The samples were washed with permeabilization wash buffer, followed by centrifugation at 400g for 2 minutes. The cell pellets were resuspended in 200μl PBS with 2% Foetal Bovine Serum (FBS) (A4766801, Gibco) and 2mM EDTA and were stored at 4°C for acquisition. Appropriate fluorescence minus one (FMO) controls were also prepared in parallel. For comprehensive details regarding this protocol refer to Gkantsinikoudi *et al., 2025*(*20*).

### FSFC data acquisition and analysis

FSFC data acquisition was performed using the Cytek® 4 laser Aurora (UV, V, B, R) spectral flow cytometer and the SpectroFlow 3.3.0 software (Cytek® Biosciences, US). For spectral unmixing, single stain reference control beads were used for all fluorescent antibodies. Heat killed cells (90°C) were used to prepare the reference control for the Live/Dead Fixable Blue stain. Multiple autofluorescence extraction was used to account for the autofluorescence signatures of different cell populations. For this, a tissue- and treatment-specific unstained sample was used.

All data was analysed using the OMIQ Software (Dotmatics, US). Cellular debris was excluded from the analysis by appropriate gating using the FSC-A vs SSC-A plot, followed by exclusion of duplets using the FSC-A vs FSC-H plot. Live cells were selected based on the Live/Dead Fixable Blue stain. Then, in the CD45 vs CD31 plot, the CD45^+^ and CD45^-^ populations were analysed separately. In the CD45^-^ gate, EC populations were analysed, while in the CD45^+^ gate, analysis of immune cell populations was performed. Both classical gating analysis and dimensionality reduction analysis was used.

For dimensionality reduction, the first step of the analysis pipeline involved data quality control, which was performed using the FlowAI algorithm(*48*). Then, manual compensation was performed where necessary. For unsupervised clustering, the FlowSOM algorithm(*49*) and the cells were clustered based on the endothelial markers CD31, TM, EMCN, LYVE1, MCAM and PDPN to generate endothelial clusters with different identities. For cluster visualisation, uniform manifold approximation and projection for dimension reduction (UMAP) plots(*50*) were used. UMAP were generated using the Euclidean metric composed of the same six EC markers with a minimum distance of 0.1, nearest neighbours of 15. Contour plots, histograms and Wishbone trajectory analysis were generated using OMIQ software as detailed(*20*).

### Image analysis and profile plot generation

For image analysis, the Volocity 3D Image analysis software (PerkinElmer, US) was used. The total area for different markers was calculated and normalised using the total tissue area or total number of nuclei. For EndMT quantification, the area of colocalization of an endothelial and a mesenchymal marker was calculated.

To visualise and quantify the distribution of different markers across the three zones of the liver tissue, profile plots were generated using Image J and the Macro plot line profile multicolour macro written by Kees Straatman, University of Leicester (Leicester, UK(*51*)), which was modified by Dr Paul Imbert. Regions of interest were selected that spanned from a portal vein (zone 1) led to a central vein (zone 3; Fig S1A). A 200 µm line was used to measure the fluorescence intensity between these two structures. The distance between the portal and central veins was expressed as a percentage and segmented into zone 1 (0-33% of line), zone 2 (34-66% of line) and zone 3 (67-100% of line) for normalisation across all samples. Area under the curve was used to calculate the fluorescence signal under the curve within each zone.

### Single cell RNA sequencing Data Acquisition (mouse carbon tetrachloride study)

Raw 10x Genomics single cell RNA sequencing data from the GSE145086 dataset were obtained from the Gene Expression Omnibus and processed in R. For each sample, the standard 10x output files (gene-cell count matrix, barcode file, and gene annotation file) were imported, and sparse matrices were constructed alongside their corresponding barcodes and gene identifiers. Gene symbols were inspected for redundancy and made unique where necessary, and each cell was assigned a unique sample specific identifier to enable accurate provenance tracking. Per cell metadata were generated for every sample, and individual datasets were converted into Seurat objects. Following quality checks to ensure alignment of barcodes, metadata, and gene features, all samples were merged into a single unified Seurat object.

For trajectory reconstruction, Seurat objects were converted into Monocle3 cell data sets using the as.cell_data_set function from SeuratWrappers. Cluster and partition information from Seurat were retained. The principal graph was learned using learn_graph, and pseudotime was estimated using order_cells, setting the endothelial cell cluster representing the quiescent state as the root. Pseudotime values were extracted and used to infer lineage progression among endothelial subpopulations. Only Monocle3’s internal pseudotime calculations were used for subsequent analysis; graphical visualization was performed in R but excluded from final interpretation.

Genes significantly upregulated in selected comparisons were converted to Entrez IDs using the bitr function in the clusterProfiler (v4.10.0) package (OrgDb = org.Mm.eg.db). Gene Ontology (GO) enrichment analysis for Biological Processes (BP) was performed using enrichGO( with the Benjamini–Hochberg adjusted p-value threshold of 0.05. The most significantly enriched biological processes were used for interpretation of functional differences among endothelial subpopulations.

All analyses were conducted in R (v4.3.3) on Apocrita HPC system (http://doi.org/10.5281/zenodo.438045). The following packages were used: Seurat (v5.0.3), Monocle3 (v1.3.7), SeuratWrappers (v0.3.1), clusterProfiler (v4.10.0), org.Mm.eg.db (v3.17.0), ggplot2 (v3.5.1), and dplyr (v1.1.4).

### Marker Gene Identification and Differential Expression

Cluster-specific marker genes were identified using Seurat’s FindAllMarkers function, retaining genes expressed in at least 25% of cells within a cluster and exhibiting a log fold change greater than 0.25. Differential expression analyses between experimental conditions were performed using FindMarkers with test.use = "MAST", and significant genes were defined by adjusted p < 0.05 and log₂FC > 0.25.

All results were exported as CSV files for reproducibility and integration with pathway enrichment analysis.

### CiteSeq sequencing Bariatric Surgery Patients

We performed antibody-augmented single cell RNA for cell populations sampled from liver tissue sampled during bariatric surgery (liver disease) or gallbladder surgery (healthy). Histological assessment of MASLD and fibrosis was conducted by a specialist pathologist according to the NASH CRN. Log-normalized TAGLN expression values were exported from the Seurat object and plotted in GraphPad Prism to compare expression between healthy and fibrotic liver samples(*52*).

### High-definition spatial transcriptomics data generation and analysis

5μm patient biopsy FFPE sections were prepared and stained according to the Visium HD FFPE Tissue Preparation protocols (GC000684, Revision D) and processed following the standard Visium HD Spatial Gene Expression Reagents Kits User Guide (GC000685, Revision D) as outlined by 10X Genomics. Prepared libraries were sequenced on the Illumina NovaSeq6000 SP lane with 100 cycles. The sequencing data was pre-processed using Space Ranger (v3.1.3, 10X Genomics), including the generation of the CLOUPE output at 8μmx8μm binning resolution. On a per-biopsy basis, endothelial bins were identified within Loupe Browser (v.8.0.0, 10X Genomics) by a Log2 Feature Sum of endothelial marker genes (PECAM1, THBD, EMCN, MCAM, LYVE1) and split into TAGLN+ and TAGLN-sub-populations (endo_TAGLN+ and endo_TAGLN-). Non-endothelial bins were defined as “rest_of_tissue”. Differential gene expression between TAGLN+ and TAGLN-endothelium-associated bins was identified within Loupe Browser using the ‘Run Differential Expression’ tool, with significance adjusted for multiple comparisons using Benjamini-Hochberg correction. To define potential receptor-ligand analyses, barcode annotations were exported from Loupe Browser. The Visium HD raw counts data matrix was loaded into a SpatialData object, subset to the relevant barcodes prior to annotation transfer, normalization and log transformation of the expression data and h5ad conversion. CellChat (v2.2.0) was used to generate a spatial CellChat object for each biopsy using the full human interaction database (CellChatDB.human) and to generate visualizations. Compute CommunProb was performed using parameters “type = truncated Mean” and “trim=0.01”.

### SteatoSITE ethics statement

Anonymised tissue was supplied with approval from the National Health Service Research Scotland (NRS) Biorepository Network (Ref: SR1032; 2 Aug 2018). Unified transparent approval for unconsented data inclusion was granted by the West of Scotland Research Ethics Committee 4 (Ref: 25/WS/0004; 16 Jan 2025), the Public Benefit and Privacy Panel for Health and Social Care (Ref: 2425-0221; 20 May 2025), institutional R&D departments, and Caldicott Guardians.

### SteatoSITE analysis

SteatoSITE v2 with follow up to end December 2023 was used. Only biopsy cases with RNAseq were used for analysis using R4.4.1. For overall survival analysis (n=505), surv_cutpoint() from the ‘survminer’ (0.5.1) with a minimum proportion 0.25 was used to define the optimal cutpoint of TAGLN expression (normalised cpm) using the maximally selected rank statistics from the ‘maxstat’ (0.7-26) package. For time-to-liver related event analysis, cases with no liver event coding prior to the biopsy date (n=435) were fit with a time-to-event proportional hazards regression model using NASH-CRN fibrosis stage, age, gender, and TAGLN expression as predictive variables using coxph from ‘survival’ (3.6-4), and the cumulative incidence function of the model at t=4000 days stratified for NASH-CRN fibrosis stage plotted as a function of TAGLN expression using a plot_surv_at_t from ‘contsurvplot’ (0.2.2).

### Statistical Analysis

All statistical analysis was completed using the GraphPad Prism 10.0.0 software (GraphPad, US). Statistical tests were performed to identify statistically significant results. The Shapiro-Wilk normality test was first used to identify whether the data are distributed normally. After that, a t-test or a one-way analysis of variance (ANOVA) was used for normally distributed data and the equivalent non-parametric test was used for data that did not pass the normality test. Appropriate multiple comparisons tests were also applied to all datasets. The exact test used for each dataset is detailed in the corresponding figure legend of each Figure. Statistical significance was defined as P≤0.05 (ns: p>0.05, *: p≤0.05, **: ≤0.01, ***: ≤0.001, ****: ≤0.0001). All data are presented as mean ± standard error of the mean (SEM).

## List of Supplementary Materials

**Supplementary Fig. 1.**
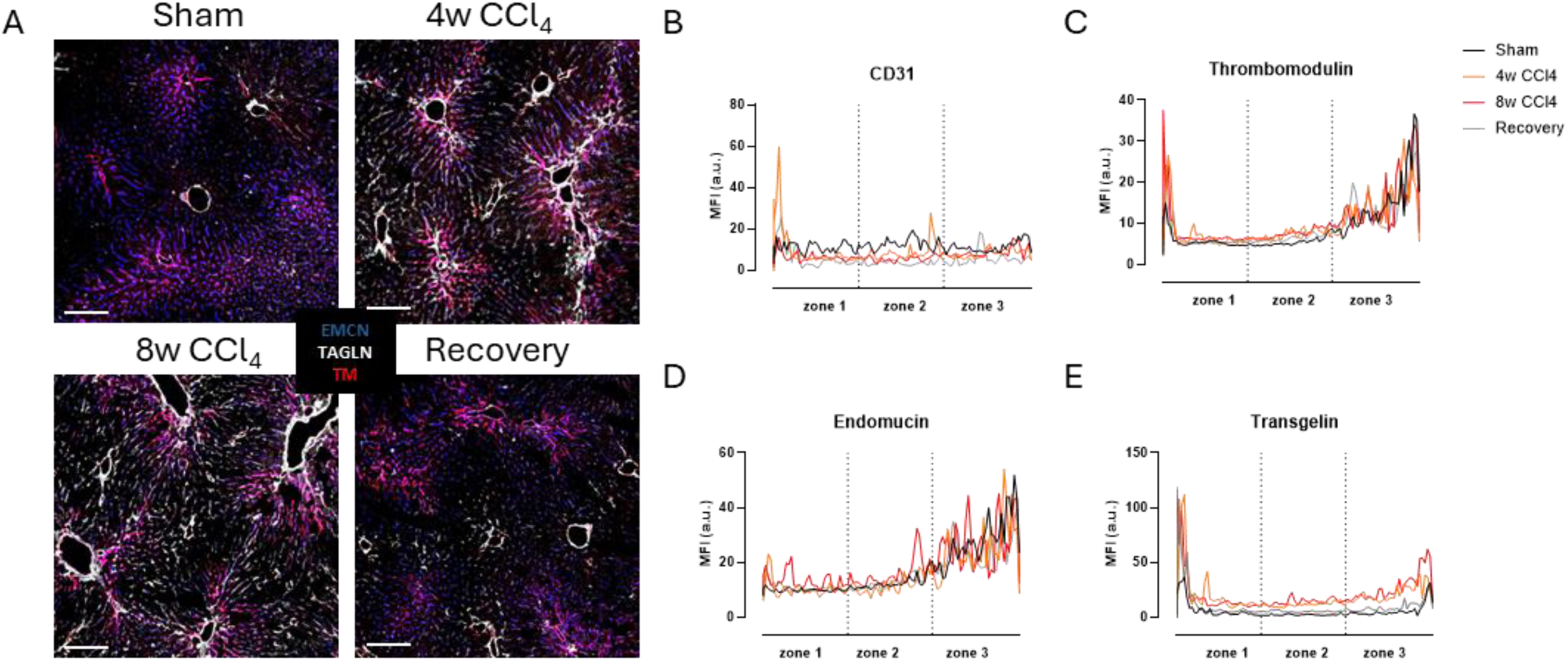
Endothelial plasticity and tissue remodelling in response to CCl_4_-induced liver fibrosis. (A) Representative confocal images of thrombomodulin (TM) (red), endomucin (EMCN) (blue) and transgelin (TAGLN) (white) staining in the liver of Sham, 4w and 8w CCl_4_-treated and Recovery group. Scale bar= 200μm. Profile plots for (B) CD31, (C) TM, (D) EMCN and (E) TAGLN showing the shift in zonation in Sham (black), 4w CCl_4_ (orange), 8w CCl_4_ (red) and recovery (grey) groups.

**Supplementary Fig. 2.**
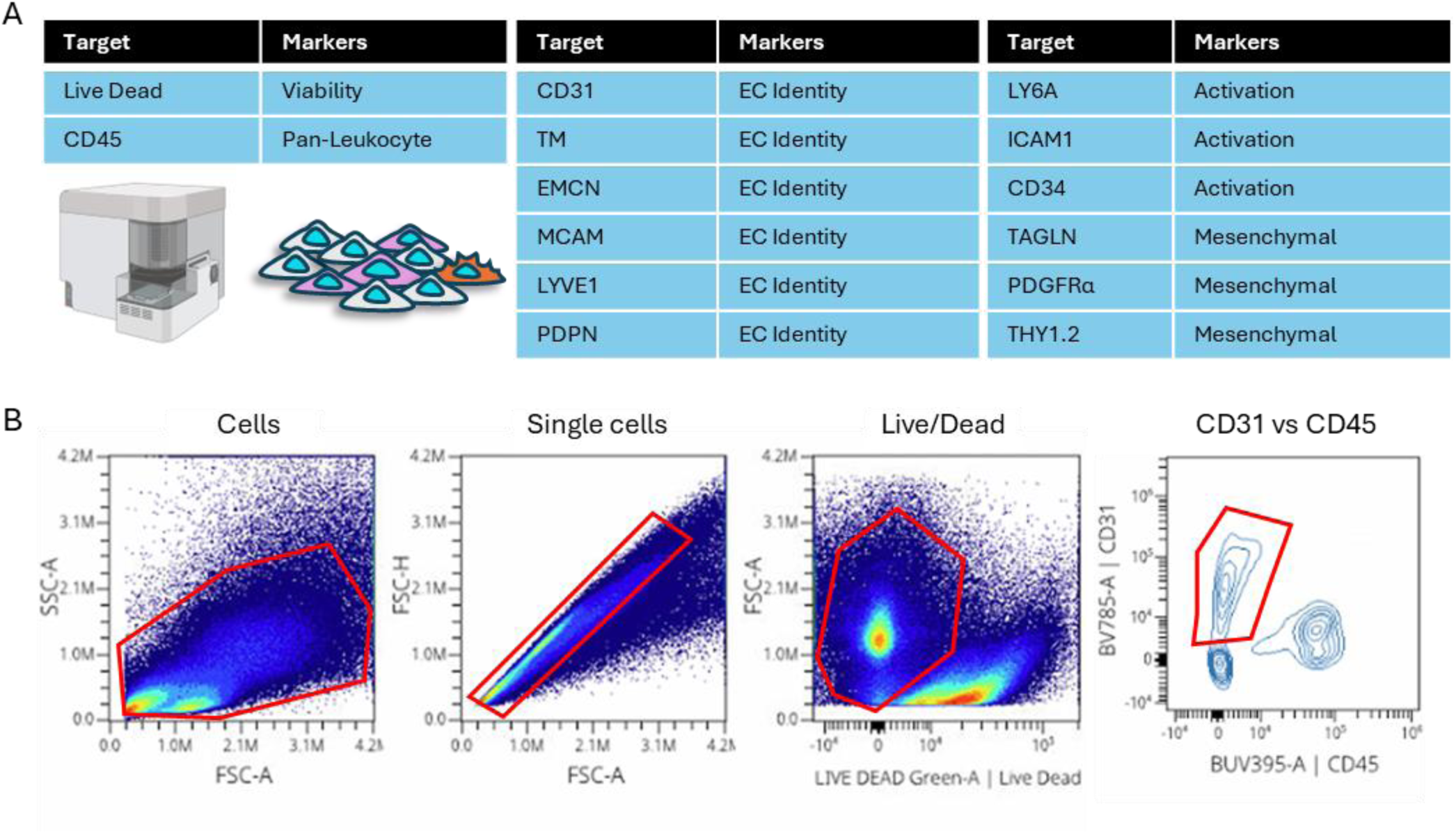
FSFC EC panel and gating. (A) Schematic for endothelial FSFC panel and (B) Gating strategy for the analysis of CD45^-^CD31^+^ endothelial cells. From left to right: a gate for all cells was firstly drawn using the forward scatter area (FSC-A) vs the side scatter area (SSC-A) plot. Then, single cells were selected using the FSC-A vs forward scatter height (FSC-H) plot. The dead cells were excluded from the analysis using the Live/Dead Fixable stain vs FSC-A plot. In the live cells gate, the CD45^-^CD31^+^ and CD45^+^CD31^-^ cells were gated separately using the CD45 vs CD31 plot. The CD45^-^CD31^+^ gate (red) was used for the analysis of endothelial cells.

**Supplementary Fig. 3.**
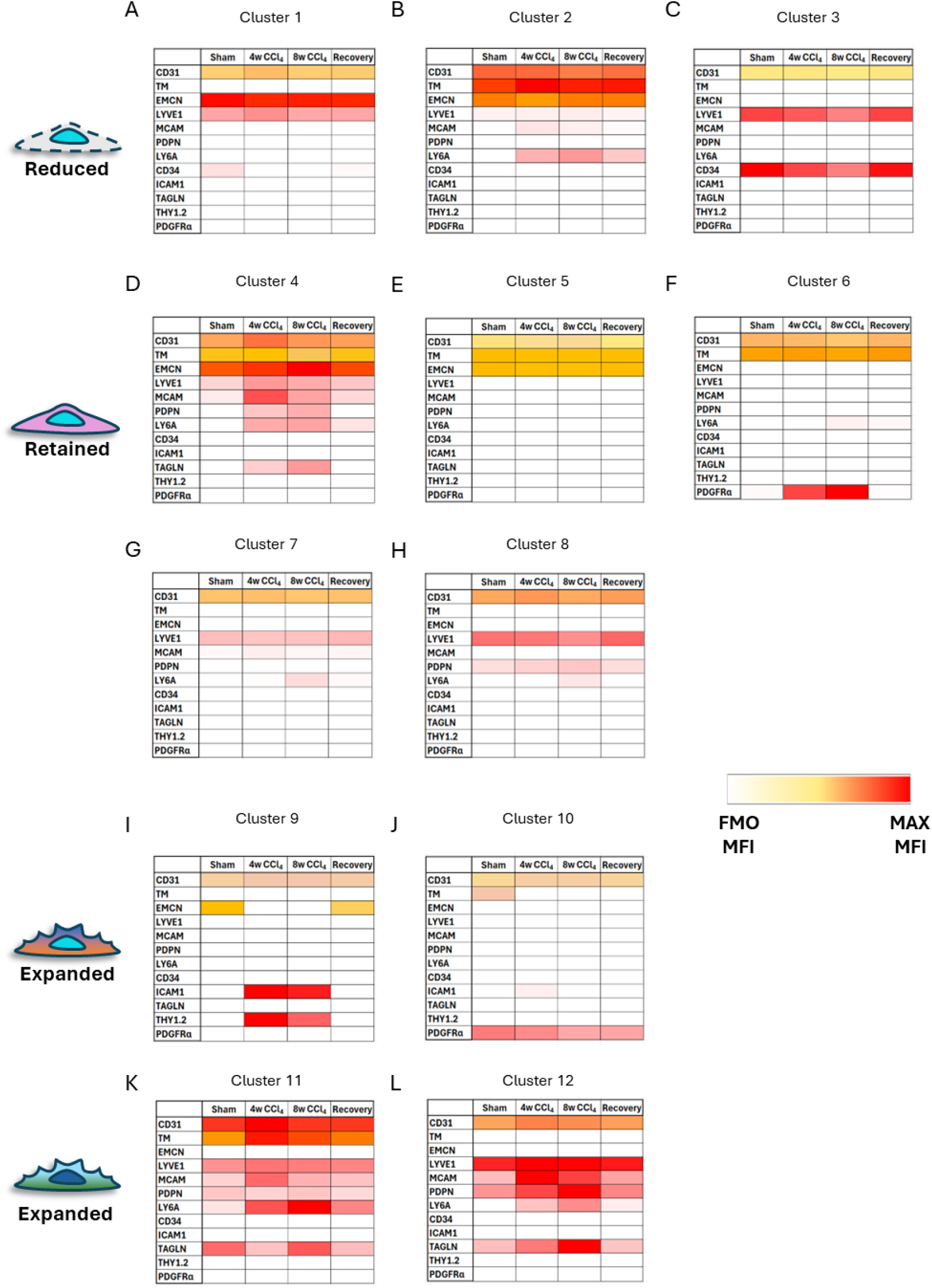
Heatmaps of the phenotypic properties of each EC cluster split by treatment group. Heatmaps of median fluorescence intensity (MFI) for (A-C) Reduced clusters 1-3 (D-H) Retained clusters 4-8 and (I-L) Expanded clusters 9-12. The heatmaps were created using full spectrum flow cytometry data. The fluorescence minus one (FMO) MFI for each marker was used as the threshold for positive signal.

**Supplementary Figure 4.**
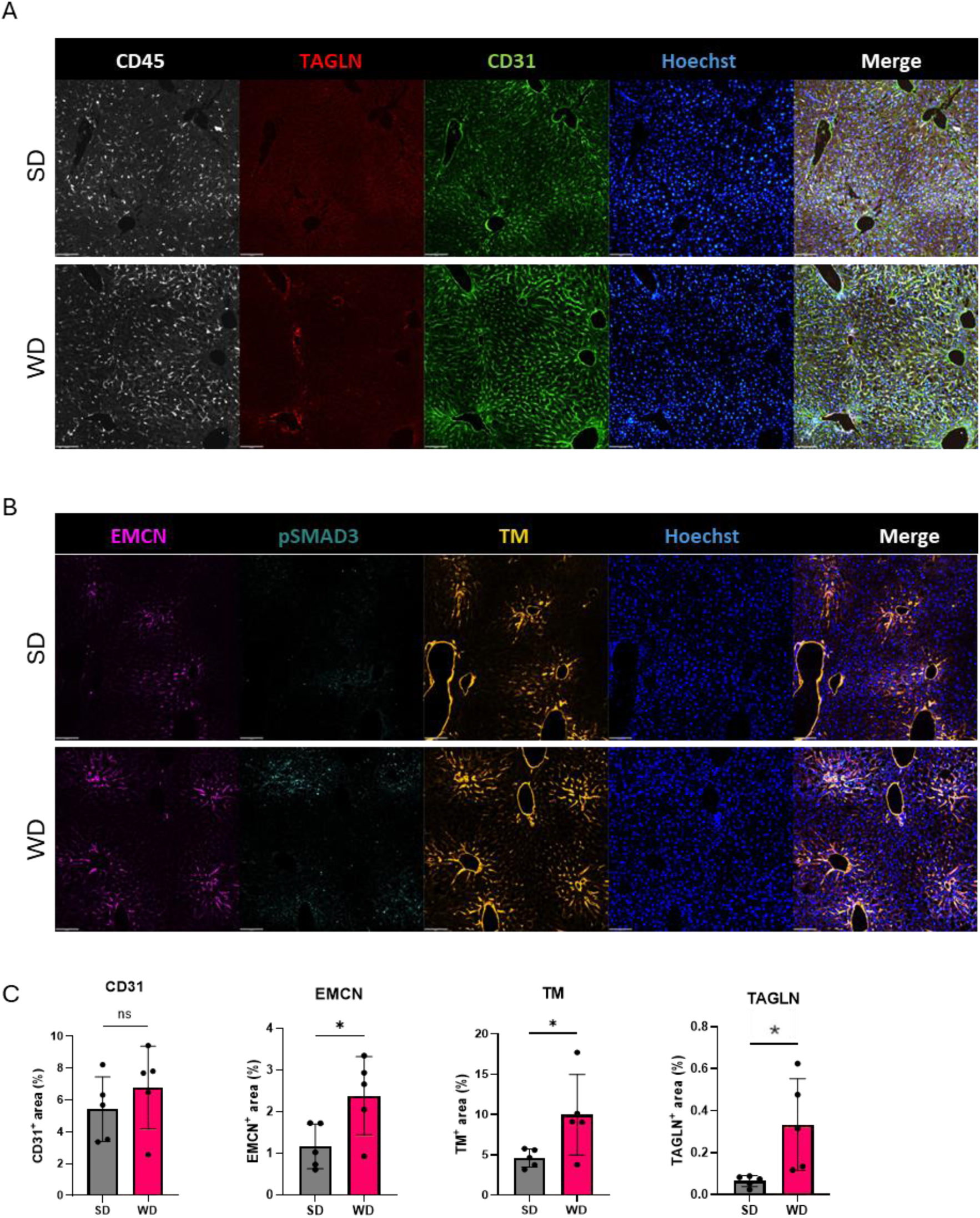
Changes in vascular patterning in response to a 12w WD. (A, B) Representative confocal IF images of liver sections from mice on a standard diet (SD) or Western diet (WD) for 12 weeks stained for (A) CD45 (white), transgelin (TAGLN) (red) and CD31 (green) and (B) endomucin (EMCN) (red), pSMAD3 (turquoise) and thrombomodulin (TM) (yellow). Scale bars= 200μm. (C) Quantification of positive fluorescence signal for (C) CD31, EMCN, TM, and TAGLN. For statistical analysis, unpaired t-tests were used. ns; not significant, *; p<0.05.

**Supplementary Figure 5.**
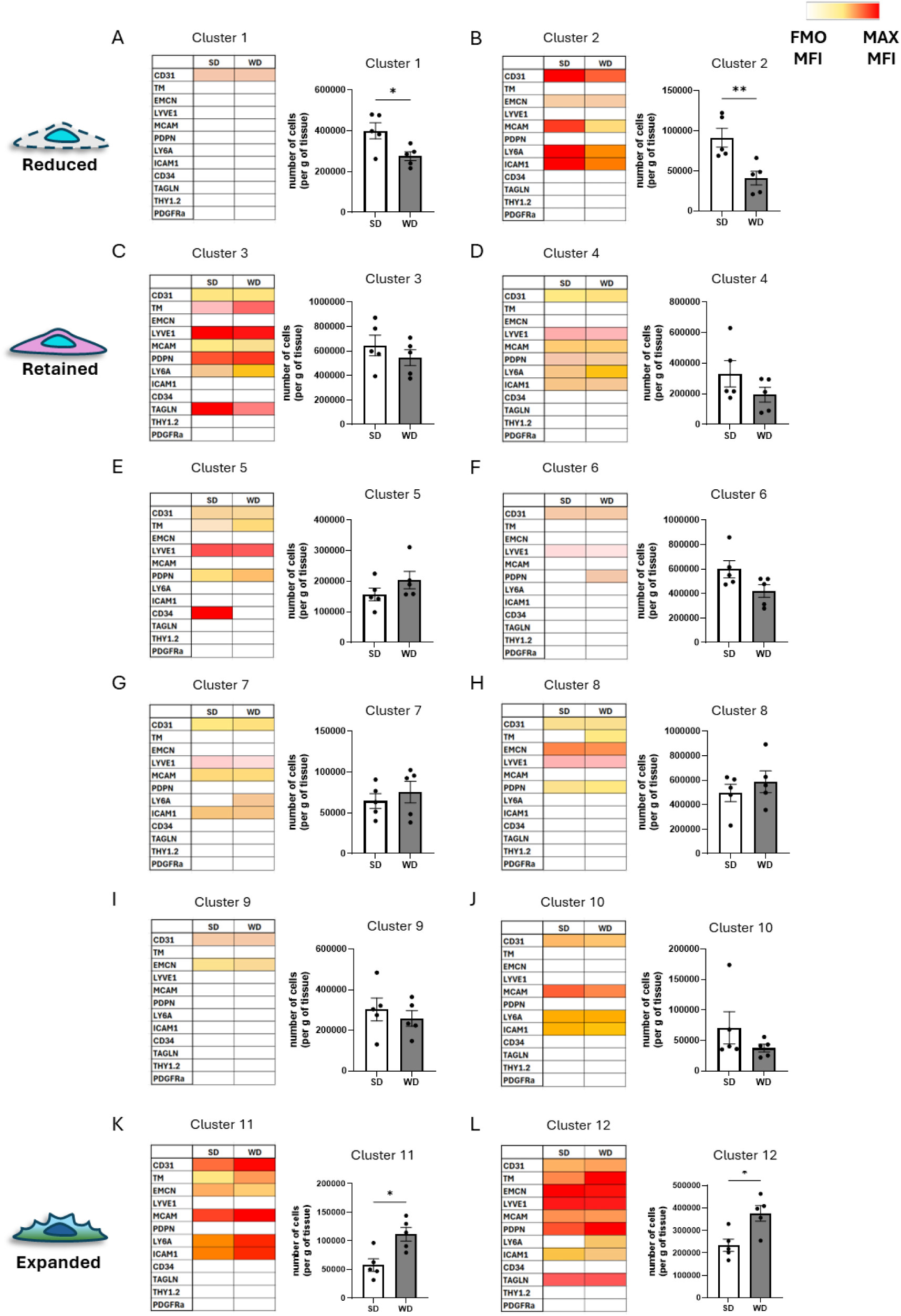
Heatmaps of the phenotypic properties of each EC cluster split by diet group. Heatmaps of median fluorescence intensity (MFI) for (A, B) Reduced clusters 1 and 2, (C-J) Retained clusters 3-10 and (K, L) Expanded clusters 11 and 12. The heatmaps were created using full spectrum flow cytometry data. The fluorescence minus one (FMO) MFI for each marker was used as the threshold for positive signal.

**Supplementary Figure 6.**
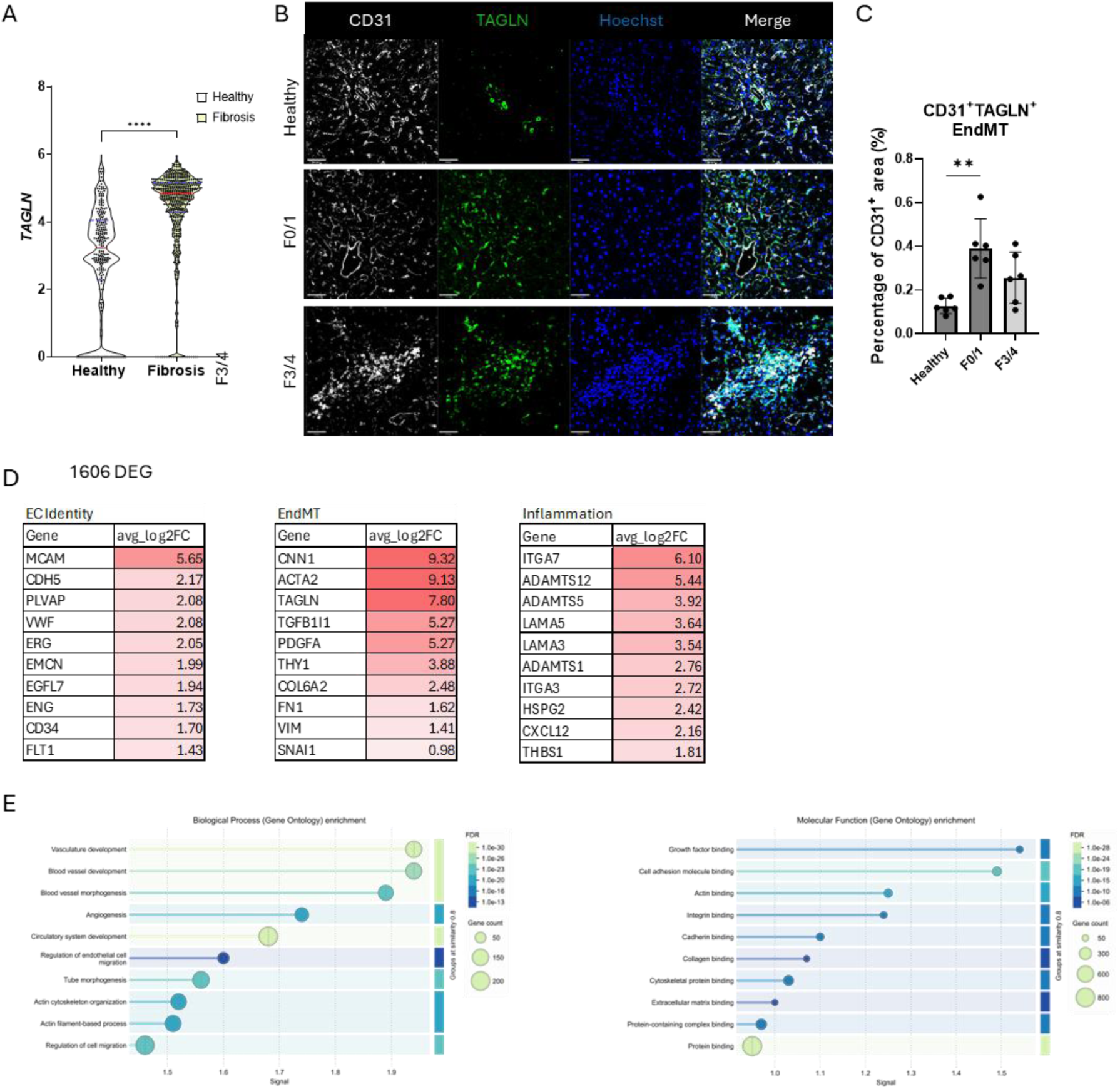
Expression of TAGLN^+^ FAEC is a conserved feature of MASLD with the potential to be a biomarker of progressive disease. (A) Violin plot of scRNAseq analysis of cell number (dots) and median expression profile of *TAGLN^+^*EC from liver biopsies from MASLD patients after bariatric surgery. (B) Representative confocal images of patient biopsies stained for CD31 (white), transgelin (TAGLN) (green) and nuclei (Hoechst; blue). Scale bar= 50 μm. (C) Quantification of CD31^+^TAGLN^+^ EndMT cells expressed as a percentage of total CD31^+^ area (N=6 patients per group). (D) Gene enrichment analysis of *TAGLN^+^* FAEC presented as a Table of top 10 enriched EC identity genes, EndMT genes and inflammation identified using scRNAseq. (E) Gene ontology analysis of all significantly enriched genes (Log2FC >0.5, P adj <0.05) for Biological Processes and Molecular Function.

**Supplementary Figure 7.**
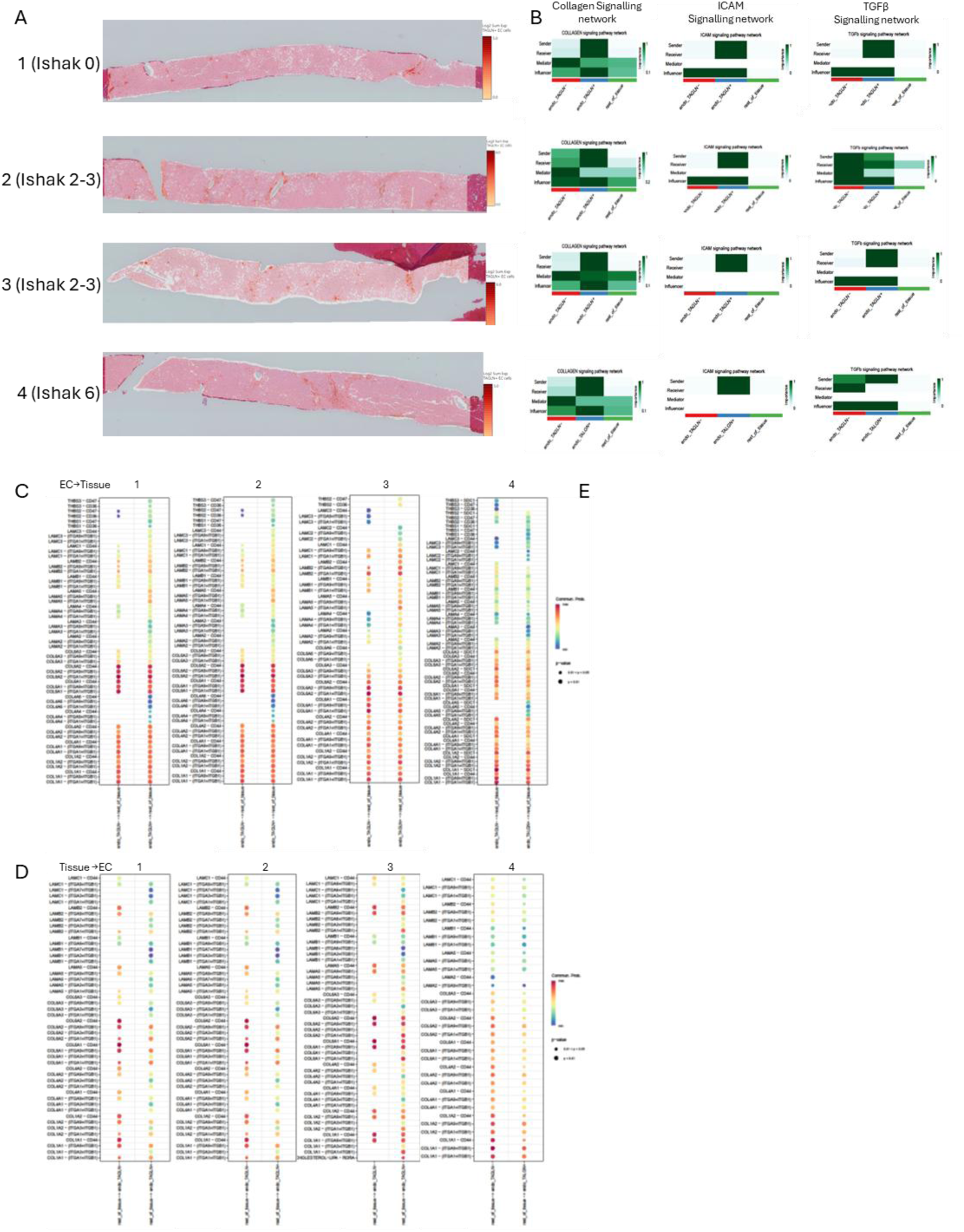
Spatial Transcriptomic analysis of four MASLD patients. (A) Localisation of TAGLN^+^EC in patients with different stratification from F0-F4. (B) Heatmap of enriched Collagen, ICAM and TGFβ signalling pathways per patient. CellChat analysis for each patient for (C) TAGLN^-^EC vs TAGLN^+^EC to tissue and (D) tissue to TAGLN^-^EC vs TAGLN^+^EC.

**Supplementary Table 1.** Patient profile for spatial transcriptomic data.

| Biopsy ID | Diagnosis | Age (years) | Ethnicity | Sex (F/M) | Ethanol (ml/ week) | BMI (kg/m<sup>2</sup>) | Liver Stiffness Measurement (kPa) | Liver Steatosis Measurement (dB/m) | Histological fibrosis stage (Ishak) | FIB-4 | Co-morbidities |
| --- | --- | --- | --- | --- | --- | --- | --- | --- | --- | --- | --- |
| 1 | MASLD | 53 | White-British | F | 170 | 48.08 | 12.6 | 265 | 0 | 0.64 | Hypertension, Type 2 Diabetes, Atrial Fibrillation, previously resected thyroid cancer |
| 2 | MASLD | 56 | White-British | M | 0 | 44.26 | 33.0 | 382 | 2-3 | 1.28 | Depression, Hypertension, Gout, Obstructive Sleep Apnoea, Osteoarthritis |
| 3 | MASLD | 21 | Mixed – White and Asian | F | 0 | 22.30 | 16.5 | 258 | 2-3 | 0.58 | Depression |
| 4 | MASLD | 48 | White-British | M | 80 | 23.13 | 15.8 | 332 | 6 | 3.37 | Nocturnal epilepsy |

**Supplementary Table 2.** List of primary (top) and secondary (bottom) antibodies used for tissue immunofluorescence experiments. (N/A: not applicable, AF: Alexa Fluor)

| Target | Host | Clonality | Fluorophore | Dilution | Product number | Supplier |
| --- | --- | --- | --- | --- | --- | --- |
| CD31 | Goat | polyclonal | AF488 | 1:100 | FAB3628G | R&D Systems |
| CD31 | Sheep | polyclonal | unconjugated | 1:100 | AF806 | R&D Systems |
| TM | Goat | polyclonal | unconjugated | 1:500 | AF3894 | R&D Systems |
| EMCN | Rat | Monoclonal (V.7C7.1) | unconjugated | 1:200 | ab106100 | Abcam |
| LYVE1 | Rat | Monoclonal (ALY7) | unconjugated | 1:100 | 48-0443-82 | Invitrogen |
| MCAM | Rat | Monoclonal (ME-9F1) | unconjugated | 1:100 | 130-101-866 | Miltenyi |
| TAGLN | Rabbit | Polyclonal | unconjugated | 1:500 | ab14106 | Abcam |
| THY1.2 | Rat | Monoclonal (G7) | unconjugated | 1:100 | 105201 | Biologend |
| phospho-SMAD3 | Rabbit | Monoclonal (EP823Y) | unconjugated | 1:100 | ab52903 | Abcam |
| CD45 | Rat | F30-11 | unconjugated | 1:100 | 103102 | Biologend |

| Host | Reactivity | Fluorophore | Dilution | Product number | Supplier |
| --- | --- | --- | --- | --- | --- |
| donkey | goat | 488 | 1:1000 | A32814 | Invitrogen |
| donkey | goat | 555 | 1:1000 | A21432 | Invitrogen |
| donkey | rabbit | 488 | 1:1000 | A21206 | Invitrogen |
| donkey | rabbit | 555 | 1:1000 | A32794 | Invitrogen |
| donkey | rabbit | 647 | 1:1000 | A31573 | Invitrogen |
| donkey | rat | 488 | 1:1000 | A21208 | Invitrogen |
| donkey | rat | 550 | 1:1000 | SA5-10027 | Invitrogen |
| chicken | rat | 647 | 1:1000 | A21472 | Invitrogen |

**Supplementary Table 3.**
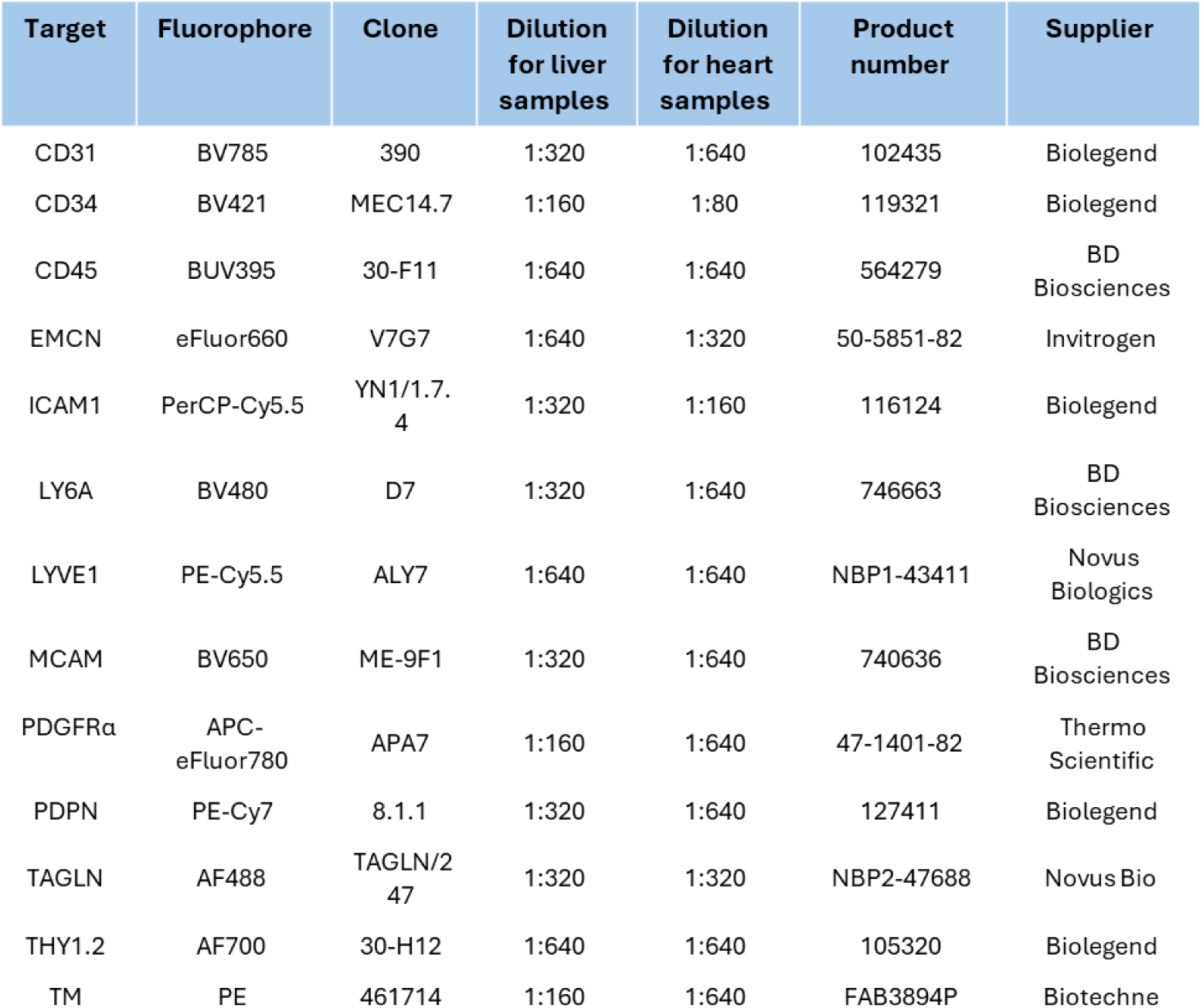
Antibody details for the endothelial panel used for full spectrum flow cytometry.

## Acknowledgements

This research was supported by Dr Paul Imbert, manager of the CMR Advanced Bioimaging Facility (CMR-ABF), and Manuela Terranova-Barberio, manager of the WHRI Flow Cytometry Facility which are both located at the William Harvey Research Institute, Charterhouse Square, Queen Mary University of London, EC1M 6BQ, United Kingdom.

Lindsay Birchall is thanked for her technical assistance in spatial transcriptomics sample preparation. The University of Manchester Genomics Technology Core Facility is thanked for their assistance with the Visium HD spatial transcriptomics protocol.

## Funding

Medical Research Council project grant MR/Y013751/1 (ND)

Bart’s Charity Seed Grant MGU0514 (ND)

British Heart Foundation PhD studentship FS/4yPhD/F/21/34161 (CG)

Medical Research Council project grant MR/T031883/1 (WA)

Innovate UK project grant TS/U00457X/1 (KPH)

Innovate UK project grant TS/R017581/1 (JAF/TJK)

Guts UK DGO2019_16 (JAF/TJK)

## Author contributions

Conceptualization: CG, JPD, AW, NPD

Methodology: CG, RK, EJ, SL, TJK, NPD

Investigation: CG, JPD, RK, EJ, TJK, WL, MS, NPD

Visualization: CG, RK, EJ, TJK, NPD

Funding acquisition: KPH, AR, WA, NPD

Project administration: AR, JAF, SL, VSL, KPH, AW, NPD

Supervision: AR, JAF, KPH, AW, NPD

Writing – original draft: CG, JPD, NPD

Writing – review & editing: CG, JPD, RK, EJ, TJK, WL, MS, AR, JAF, KPH, AW, NPD

## Data and materials availability

Murine scRNAseq data were analyzed from 10x Genomics single-cell transcriptomic data of isolated non-parenchymal liver cells from healthy and diseased (2w and 4w CCl4 treated) mice deposited on the GEO database (GSE145086).

For access to the full SteatoSITE resource, including rich clinical outcome annotation please visit https://regeneration-repair.ed.ac.uk/research/research-resources/steatosite. Raw RNA-seq data (FASTQ files) and minimal metadata (fibrosis stage) is available in the European Nucleotide Archive (https://www.ebi.ac.uk/ena; study accession number: PRJEB58625).

Data from the Citeseq (Fig. S6) and Visium HD Spatial Transcriptomics analyses (Fig. 8 and Fig. S7) will be made available on the GEO databases upon acceptance.

